# Sequence-based correction of barcode bias in massively parallel reporter assays

**DOI:** 10.1101/2021.04.29.442047

**Authors:** Dongwon Lee, Ashish Kapoor, Changhee Lee, Michael Mudgett, Michael A. Beer, Aravinda Chakravarti

**Author notes:** Send all correspondence to: Dongwon Lee, Ph.D., Division of Nephrology, Boston Children’s Hospital & Harvard Medical School, 300 Longwood Ave, Enders 505, Boston, MA 02115, USA, (T), (E). Division of Nephrology, Boston Children’s Hospital, Boston, MA, USA.

## Abstract

Massively parallel reporter assays (MPRA) are a high-throughput method for evaluating *in vitro* activities of thousands of candidate cis-regulatory elements (CREs). In these assays, candidate sequences are cloned upstream or downstream of a reporter gene tagged by unique DNA sequences. However, tag sequences may themselves affect reporter gene expression and lead to major potential biases in the measured cis-regulatory activity. Here, we present a sequence-based method for correcting tag sequence-specific effects and demonstrate that our method can significantly reduce this source of variation, and improve the identification of functional regulatory variants by MPRAs. We also show that our model captures sequence features associated with post-transcriptional regulation of mRNA. Thus, this new method helps to not only improve detection of regulatory signals in MPRA experiments but also to design better MPRA protocols.

## INTRODUCTION

Functional characterization of cis-regulatory elements (CREs) and their sequence variants is an essential but challenging first step for understanding gene expression regulation and how sequence variation impacts this process. Given the large expected number of regulatory variants in the human genome, and their pivotal role in human complex traits (Maurano et al. 2012; Albert and Kruglyak 2015; Lee et al. 2018), there is a pressing need for experimental methods that can rapidly test elements for cis-regulatory activity to identify CREs and their variants en masse for detecting causal effects. Massively parallel reporter assays (MPRAs) (Patwardhan et al. 2009, 2012; Melnikov et al. 2012; Kwasnieski et al. 2012) are promising for this purpose as studies identifying functional regulatory variants underlying complex human disease and traits have shown (Tewhey et al. 2016; Ulirsch et al. 2016).

Several different designs have been proposed for MPRA experiments. In a typical MPRA, CREs of interest are cloned upstream of a minimal promoter-driven reporter gene uniquely tagged by short DNA sequences (tags) placed at the 3’ untranslated region (UTR). Different locations of tags and CREs have also been explored. Self-transcribing active regulatory region sequencing (STARR-seq) (Arnold et al. 2013) places CREs downstream of the reporter’s transcript start site and uses the transcribed CREs as “tags.” More recently, some of the MPRA designs place tags in the 5’ UTR of the reporter (Klein et al. 2020). CRE activities are then measured by relative read counts of the transcribed tags versus the input library tags, using high-throughput sequencing. These analyses are, however, subject to potential biases in the detected reporter expression originating from tag sequence-specific effects. Prior studies (Melnikov et al. 2012; Ulirsch et al. 2016) have clearly shown reproducible effects of tag sequence on reporter readout, suggesting sequence-specific effects on expression, likely from mRNA stability and/or RNA-binding proteins, but efforts to identify sequence features responsible for such biases have not been very satisfactory (Melnikov et al. 2012; Ernst et al. 2016). To alleviate this problem, most MPRA studies average the effects of multiple tags (≥10) per CRE. This tactic, however, does not remove the root problem because the biases are sequence-dependent and non-random, and tag sequences are not uniformly distributed across CREs. Ideally, a model that explains the effect of tags in DNA sequence level would not only refine the MPRA results but also help to design better MPRAs. To address this issue, we introduce here a sequence-based, machine learning method, <u>M</u>PRA <u>T</u>ag <u>S</u>equence <u>A</u>nalysis (MTSA), for predicting and correcting tag sequence-specific effects.

## RESULTS

### Tag sequence significantly contributes to expression variation

Our primary hypothesis is that variation in tag expression assigned to the same CRE can be predicted from tag DNA sequence: our strategy is outlined in **Fig. 1** (see **Methods**). As proof of principle, we analyzed 20 publicly available MPRA data sets from nine different studies (Melnikov et al. 2012; Ulirsch et al. 2016; Kheradpour et al. 2013; Inoue et al. 2017; Ernst et al. 2016; Mogno et al. 2013; Kwasnieski et al. 2014; Klein et al. 2020) in which individual tag level expression and their sequences were available (**Supplemental Table S1**). Briefly, we first calculated relative tag expression, i.e., tag expression normalized to a zero mean across all tags in one set associated with the same CRE, to eliminate CRE-to-CRE variation in expression. Tags with small read counts from input libraries exhibit much larger variation in their relative expression (**Supplemental Fig. S1**). Since this variation is likely caused by random sampling and PCR amplification of low representation tags in MPRA libraries, we excluded them from model training and subsequent analyses. In fact, removal of such less-abundant tags is already common practice in MPRA studies.(Ulirsch et al. 2016; Kheradpour et al. 2013; Inoue et al. 2017; Kwasnieski et al. 2014). We note that tag-sequence-specific biases of these excluded tags can still be corrected after building models. Next, we trained support vector regression (SVR) models (Drucker et al. 1997) to learn the contribution of each tag sequence to its relative expression. We used gapped *k*-mers as sequence features (Ghandi et al. 2014) and developed new software based on LS-GKM (Lee 2016) and the SVR routines implemented in LIBSVM (Chang and Lin 2011). The trained SVR models achieved high Pearson correlations between observed and predicted relative expression (r = 0.4~0.7), with 5-fold cross-validation for most data sets (Fig. 2A; **Supplemental Fig. S2**). These results confirm that a significant fraction (up to 50%) of expression differences between tags corresponding to a specific CRE arise from tag sequence effects. Interestingly, MPRA data sets from Klein’s study (Klein et al. 2020) showed a varying degree of correlations between the observed and predicted expression (r = 0.08 ~0.57) (**Supplemental Fig. S3**). While we achieved reasonable correlation for STARR-seq libraries, we consistently had less correlation for the other types of MPRA designs. Since all the other libraries are derived from STARR-seq libraries in Klein’s study, we speculate that additional cloning steps to construct different MPRA designs may have introduced technical biases not related to the tag sequences.

**Figure 1:**
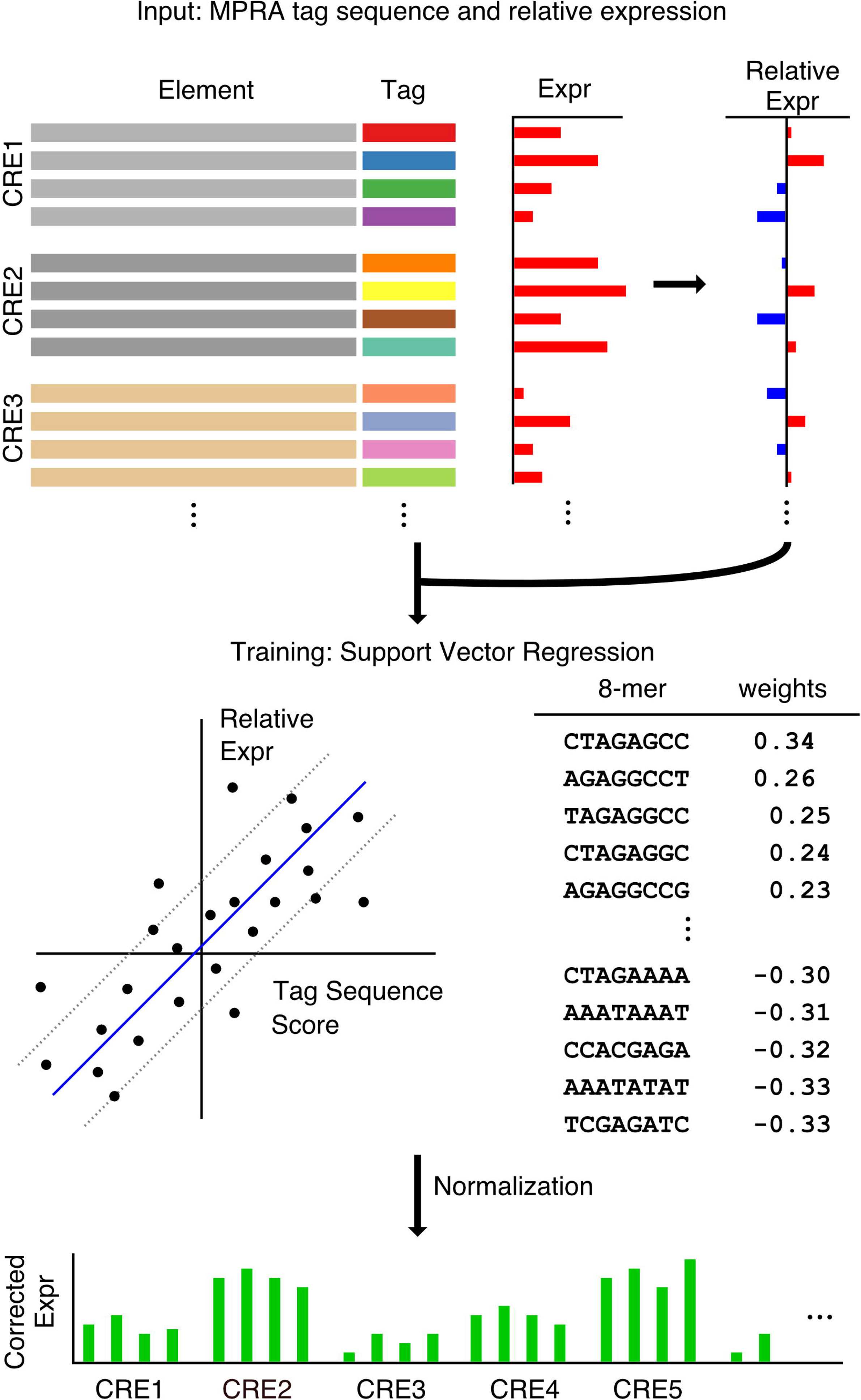
Overview of MPRA tag sequence analysis. For each tag, relative expression within each of the CREs is calculated. Tag sequences and their relative expression are then used to train Support Vector Regression (SVR) models using gapped *k*-mers as features. Tag sequence effects on their expression, as learned by SVR, are used to correct biases in raw expression data.

**Figure 2:**
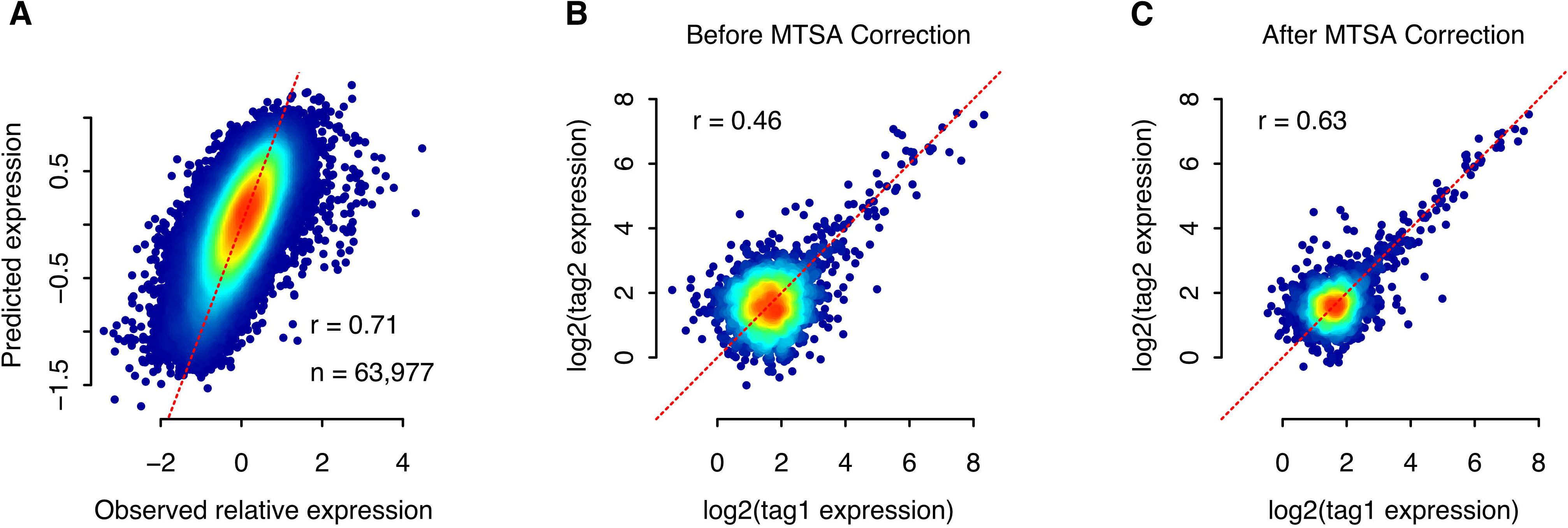
MTSA accurately predicts tag sequence effects on expression. **(A)** Observed relative expression is compared to its predicted value from tag sequence with 5-fold cross validation using Ulirsch’s MPRA data. **(B, C)** MTSA correction improves the correlation between two randomly selected tags from each CRE: log_2_ transformed raw expression **(B)** and MTSA corrected expression **(C)** are compared.

We note that MTSA uses two rounds of training to build more accurate models. The first round with 5-fold cross-validation corrects CRE-wide bias (i.e., CREs associated with tags that only increase or decrease expression). Subsequently, this corrected expression is used in the second round to build the tag sequence-specific model. We discovered that this approach consistently improved the correlations between the observed and predicted expression by 0.01 - 0.14 in all cases (**Supplemental Table S2**). These results also confirm that simple averaging will not eliminate tag sequence-dependent biases, especially if a small number of tags per CRE is used. Also, in contrast with our previous experience with building sequence-based models for CREs (Lee et al. 2011), the use of reverse-complement sequences as distinct features marginally but consistently improved model performance (**Supplemental Table S3**). This result implies that the effect exerted by transcribed tags (single-strand RNA), which distinguish reverse complement *k*-mers, is more influential on reporter expression than the effect exerted by non-transcribed tags (double-strand DNA) which do not distinguish such *k*-mers. Thus, we hypothesize that the major molecular mechanisms by which tag sequences affect reporter expression are post-transcriptional, involving mRNA stability and processing, but not transcriptional, involving CREs. To further demonstrate that MTSA can reduce sequence-specific biases in absolute expression, we compared raw expression between two randomly selected tags from each CRE, before and after MTSA. To avoid overfitting, tags used for comparison were excluded before training. Expectedly, the correlation of MTSA-corrected expression between tags is consistently higher than the original values (**Fig. 2B, C**; **Supplemental Fig. S4**).

### Tag-sequence-based correction of MPRA expression improves the precision of reporter activity measurement

MTSA corrections provide enhanced statistical power for identifying functional CRE variants by reducing tag-to-tag expression variation unrelated with the CRE effect, although it marginally affects CRE expression. To demonstrate this, we first compared CRE-level expression and their standard deviation between tags before and after MTSA correction. For most data sets we tested, correlations of CRE-level expression before and after MTSA correction are high (r > 0.95), and the variance between tags is significantly reduced after correction (**Supplemental Fig. S5**). We also tested how the number of tags affects MTSA correction. Using the Ino17 data where 100 tags per CRE were available, we down-sampled the data to simulate conditions with fewer tags per CREs. We then compared their CRE-level expression to those with all 100 tags, with and without MTSA correction. The MTSA correction consistently improved the correlation, especially when the number of tags were < 50 (**Supplemental Fig. S6**). On the other hand, our model does not improve the correlation of CRE-expression between experimental replicates because they have an identical set of tags (**Supplemental Table S4**).

Since the effects of single DNA variants on gene expression are typically small to modest, even small improvements allow improved identification of functional variants. To illustrate the power of such improvement, we applied these methods to a MPRA data set designed to find human regulatory variants affecting human red blood cell traits (Ulirsch et al. 2016). Using the raw data, we confirmed 33 (54%) of 61 variants and found 4 new significant SNPs after MTSA correction (**Supplemental Notes**). Further, variants that became statistically insignificant after MTSA correction (*n* = 28 of 61) were predicted to have a much smaller impact on CRE activity by deltaSVM, a machine-learning method for predicting regulatory effects from sequence substitutions (Lee et al. 2015), than those that remained significant (*n* = 33 of 61; Mann-Whitney *U* test, P=0.001). Moreover, the proportion of SNPs whose direction of effects agreed with their deltaSVM-predicted direction also increased from 66% (40 of 61) to 78% (29 of 37) after MTSA correction. Thus, MTSA correction improved the statistical power of the analyses considerably (**Fig. 3**).

**Figure 3.**
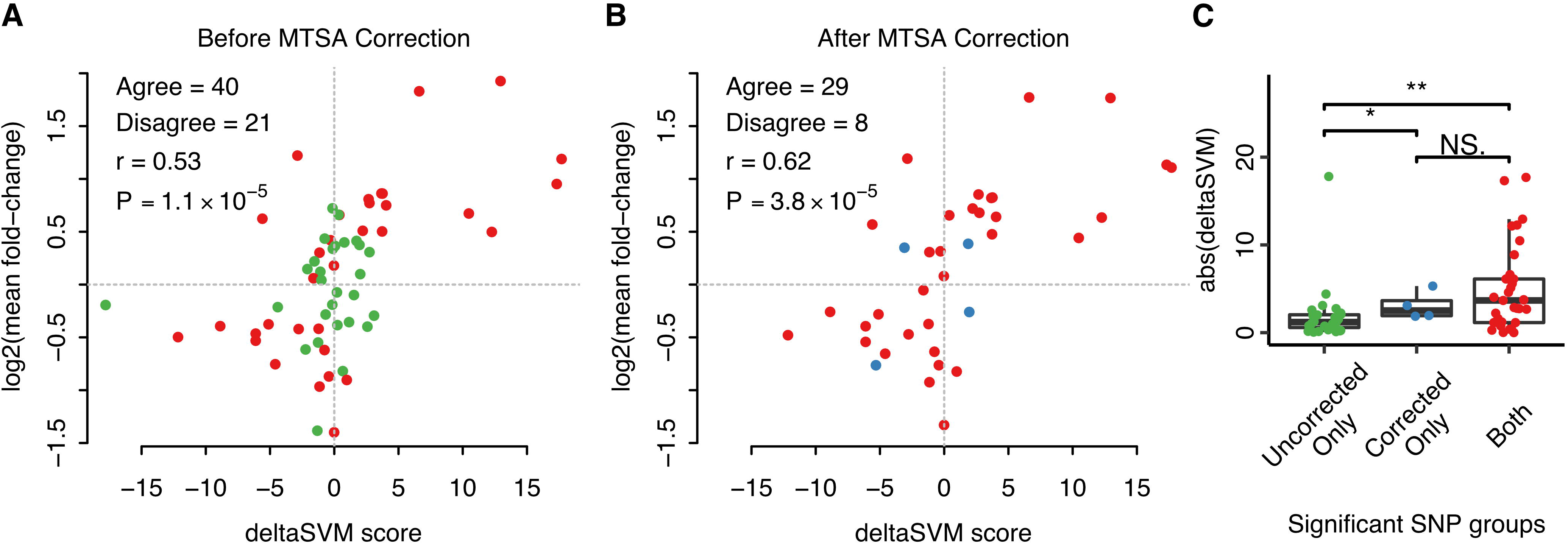
MTSA correction improves the identification of regulatory variants. **(A, B)** MTSA correction not only significantly reduces false positive identification but improves the correlation between deltaSVM and MPRA expression (from 0.53 to 0.62): log_2_ fold changes using raw expression **(A)** and MTSA corrected expression **(B)** are compared to their deltaSVM scores. SNPs detected by both are highlighted in red. **(C)** Predicted impact on CRE activity by deltaSVM is compared by group: significant SNPs without MTSA correction only (green), significant SNPs with MTSA correction only (blue), significant SNPs both with and without MTSA correction (red). The significant SNPs after MTSA correction (blue and red) are more explainable by their sequence changes (*i.e.*, larger absolute deltaSVM values). One-sided Mann-Whitney U test results are shown (*P<0.05; **P<0.005).

### Tag sequence effect on expression depends on both MPRA designs and cellular context

MTSA training assigns a weight to each of the *k*-mers, which can be interpreted as its relative contribution to expression. The weights can also be used as a scale to compare different data sets even if they don’t have the same set of tags. Pairwise comparisons using these weights uncover interesting aspects of tag effects. We found that both the experimental design and cellular context are important factors (**Fig. 4**). Seven data sets (Mel12, Khe13, Khe13K, Ern16, Ern16K, Uli16, Uli16G) use the same design initially introduced by Melnikov (Melnikov et al. 2012). These studies can be further divided into two subgroups: four studies utilizing the K562 cell lines (Khe13K, Ern16K, Uli16, and Uli16G), and the other three (two HepG2 [Khe13, Ern16], one HEK293T cell line [Mel12] studies). These two groups exhibit strong intra-group correlation with no or weak correlation in between. Moreover, there is essentially no correlation between Khe13 vs. Khe13K, as well as Ern16 vs. Ern16K that share the same MPRA design, suggesting that cellular context can strongly influence the tag sequence effect in MPRA studies. Conversely, MPRA designs also strongly influence the tag-sequence effects; for example, the SVR weights from six data sets from HepG2 cell lines (Khe13, Ern16, Ino17, Ino17W, HSS, and ORI) were only strongly correlated when the MPRA design was identical. In sum, both experimental designs and cellular contexts are major factors determining the sequence-specific effect of tags.

**Figure 4:**
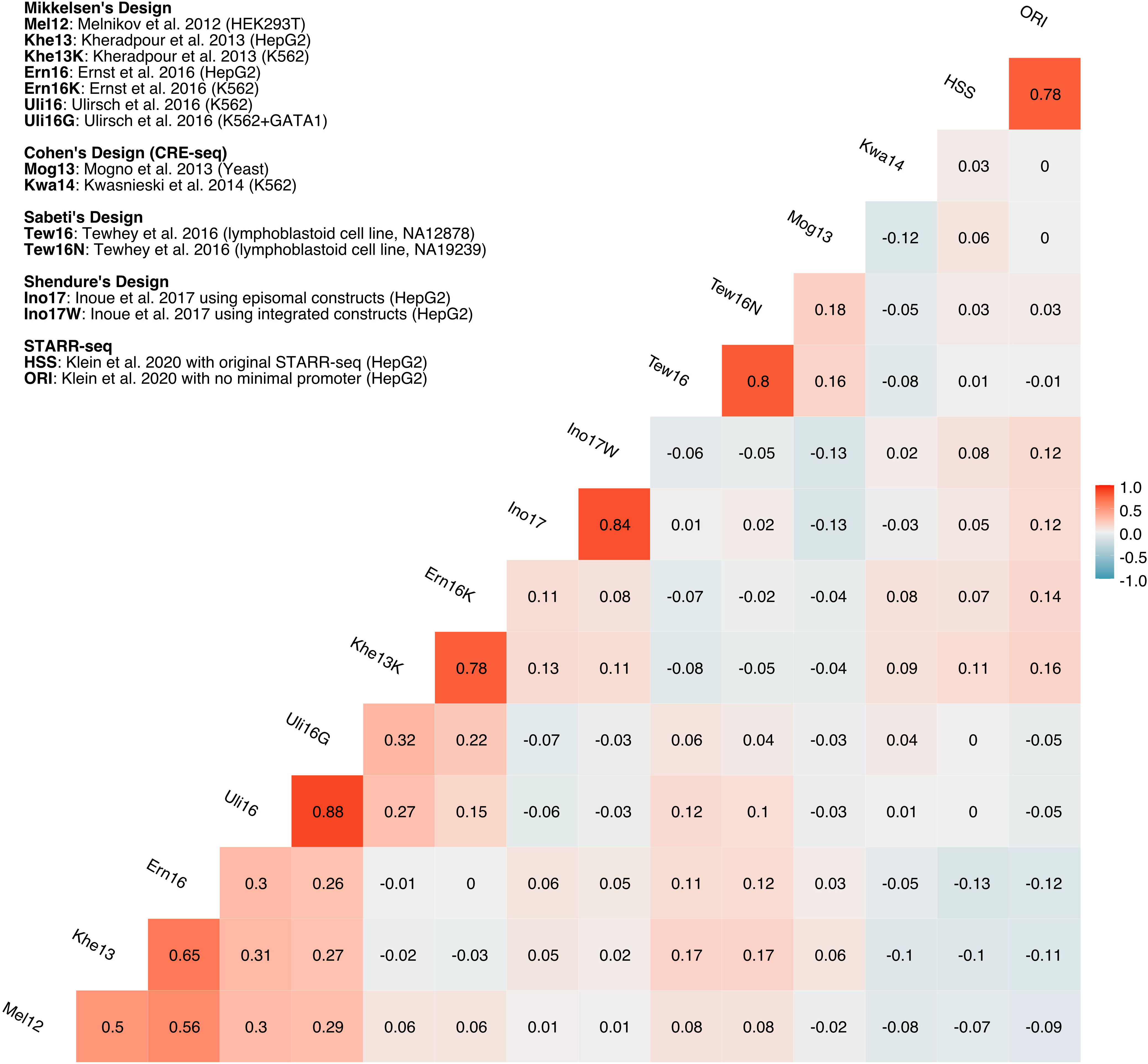
Pairwise Pearson correlations between the 15 data sets using trained 8-mer SVR weights. Differences in the MPRA designs and cell types are major determining factors of the tag sequence effects. Pearson correlation coefficients are shown in the matrix.

### Post-transcriptional regulation influences MPRA expression through tag sequences

Tag sequence effects can arise from several potential molecular mechanisms such as 3’UTR effects on gene expression (Mignone et al. 2002; He and Hannon 2004; Dominguez et al. 2018). Our approach now enables us to perform quantitative investigations of these molecular mechanisms and provides a basis for their experimental validation. Specifically, we considered five distinct biological features that could influence MPRA expression: (1) base (A, C, G, or T) counts in tag sequences (2) CREs, (3) miRNA binding sites, (4) RNB Binding Protein (RBP) binding sites, and (5) reflected stability of mRNA secondary structures. We computationally estimated the effect of each of these features from tag sequences and estimated their contribution to the tag sequence effect on reporter expression (**Methods**). Briefly, we used sequence-based prediction models (gkm-SVM)(Ghandi et al. 2014; Lee 2016) trained on open chromatin regions (DNase-seq) from the corresponding cell lines to predict the effect of tag sequences as CREs. Regarding miRNA binding sites, we only evaluated the most highly expressed miRNAs (n=20) since miRNAs with lower expression are rarely active, as previously shown (Mullokandov et al. 2012). Similar to miRNA, we considered highly expressed RBPs only and scanned the RBP binding sites using motifs from CISBP-RNA database (Ray et al. 2013). To predict the stability of the mRNA secondary structures, we padded tags with their flanking sequences (±50bp) and estimated their minimum free energy (MFE) using Vienna RNA software (Gruber et al. 2008). Next, we built multivariate linear regression models and compared their adjusted r^2^. We found that 14~51% of tag sequence-specific effects can be attributed to them, among which base counts and RBP binding sites are the most significant factors as these two feature sets explain >90% of the signals explained by all five feature sets combined (**Fig. 5**). As a negative control, we repeated the same analyses using randomized tag sequences and found that they only explained <1% of the variance (**Supplemental Fig. S7**), further supporting the significance of our results.

**Figure 5.**
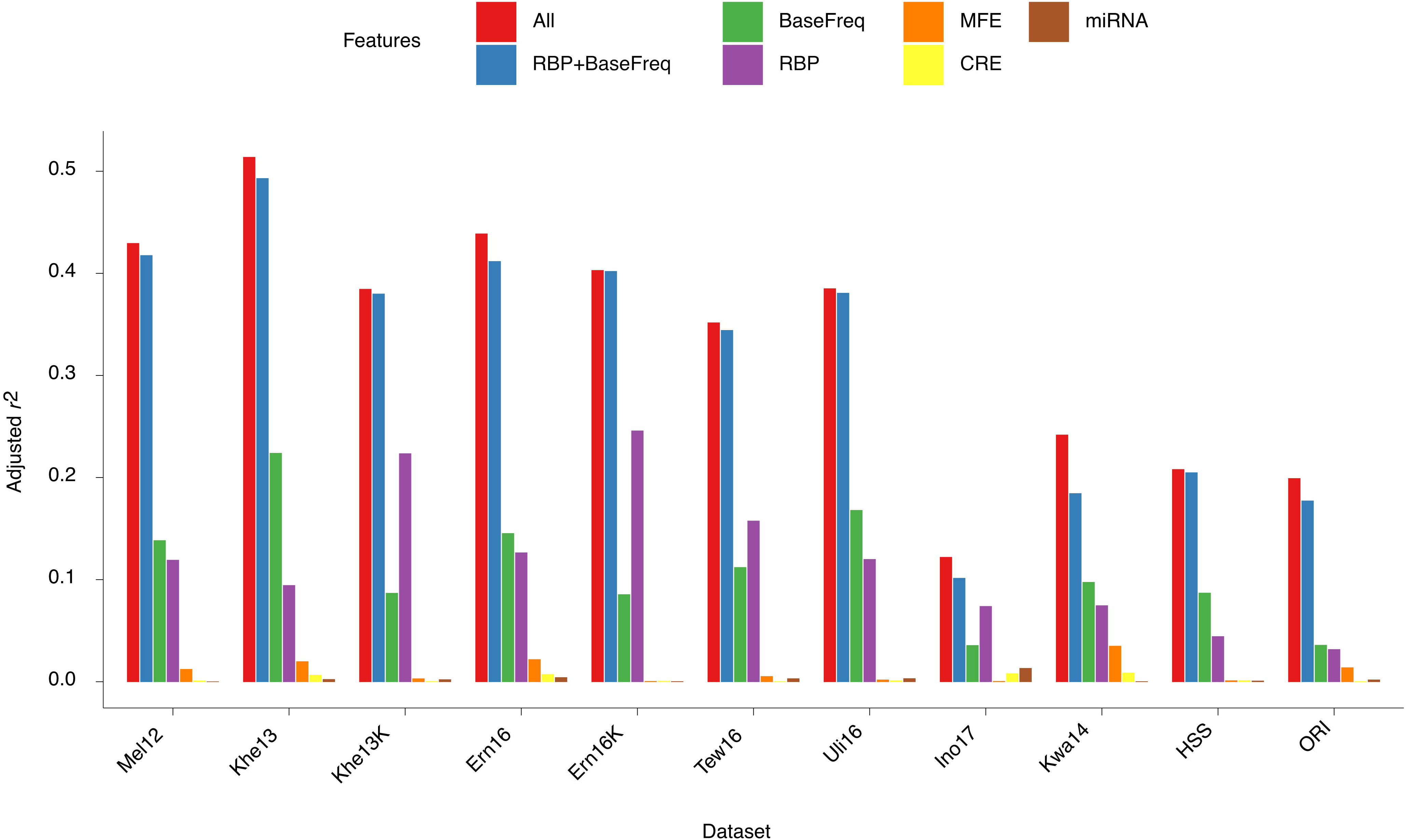
Variation in relative tag expression is explained in part by RBP binding sites and base frequencies in tags. Multivariate linear regression analyses were performed using five different biological feature sets. We used residuals of predicted expression for the individual feature set analysis after correcting for other features. BaseFreq and RBP are the two most dominant feature sets. **BaseFreq:** base counts in tag sequences, **MFE:** minimum free energy of mRNA secondary structures, **CRE:** tags as cis-regulatory elements, **miRNA:** microRNA binding sites, **RBP:** RNA binding protein binding sites, **RBP+BaseFreq:** RBP and BaseFreq features combined, **All:** all features combined.

We also evaluated these two feature sets in more detail to gain insights into potential biological mechanisms. We first assessed the base counts since multiple molecular mechanisms can be associated with them. We directly compared the base counts and the relative tag expression after adjusting other features (MFE, CRE, miRNA, and RBP). We found that the base count effects are highly specific to experiment designs and cell types (**Supplemental Figure S8**). For example, while base ‘T’ is negatively associated with the expression in Tew16, Ern16, and Mel12, it is positively associated with HSS, Khe13K, Ino17, and Kwa14. We also found that the base counts significantly correlate with the estimated minimum free energy of mRNA secondary structure (**Supplemental Fig. S9**). Thus, if non-adjusted tag expression was used, MFE also becomes strongly associated with tag-sequence specific expression for several data sets (**Supplemental Fig. 10**), suggesting that mRNA stability can be another potential molecular determinant. Next, we systematically compared effect sizes of RBP binding sites from our regression models across the data sets (**Fig. 6**). We first discovered that many RBP binding sites are present in a study-specific way due to a significant coverage difference in tag sequences. For the RBP binding sites present in multiple studies, the direction of effect is largely concordant, but its size is cell-type specific. For example, SNRPA (M348_0.6) shows a strong negative effect in K562 (Uli16, Khe13K, Ern16K), but not in the other cell types, which is consistent with its expression pattern. Lastly, we extracted the ten most positively and negatively associated 8-mers from each of the data sets and compared them with RBPs (**Supplemental Table S5**). We found that a few strong RBPs, such as SNRPA (M348_0.6), match some of these 8-mers (**Supplemental Table S5**). Intriguingly, many of these high-scoring 8-mers contain flanking sequences of the tags. One interesting case is the right flanking sequence of Melnikov’s design, AGATC. It is not only enriched in these top-scoring 8-mers, but also partially matches the SNRPA (M348_0.6) binding site, which has the strongest effect on the tag expression in K562. This result is consistent with the fact that adding flanking sequences improves the model performance significantly, especially for the K562 data sets, such as Khe13K and Ern16K (**Supplemental Table S6**).

**Figure 6.**
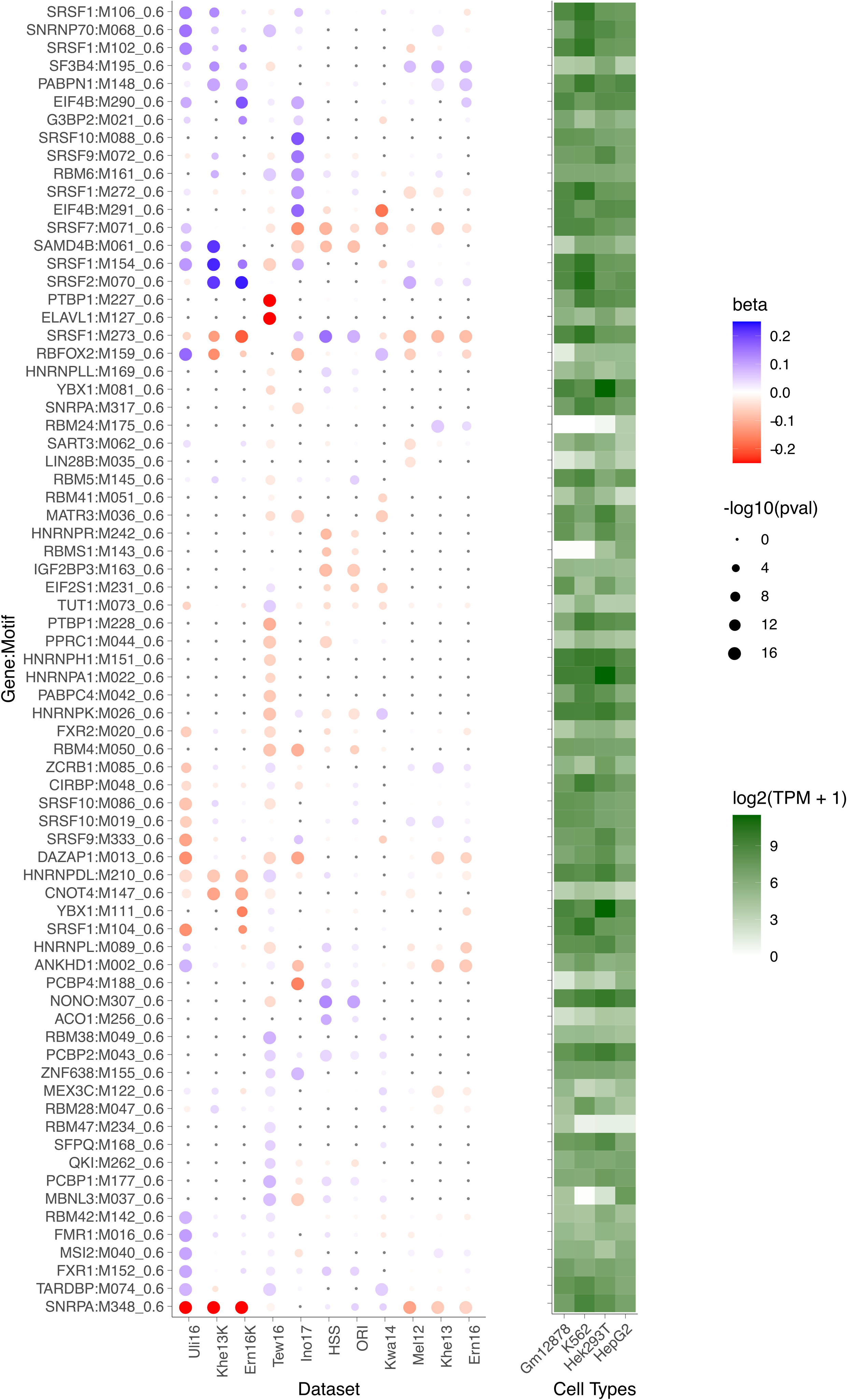
Effects of RBP binding sites in tags on relative expression is cell-type specific. **(Left)** Estimated effect sizes of RBPs from the multivariate linear regression model are compared across data sets. Only the RBPs with p-value < 10^−6^ in at least one data set are considered. RBPs that were not evaluated in the corresponding data set are shown as gray circles. **(Right)** Expression level of RBPs as log_2_(TPM + 1) in the four cell lines.

## DISCUSSION

MTSA is, to the best of our knowledge, the first method for predicting and correcting tag sequence effects on reporter gene expression, using tag DNA sequence. We expect that MTSA will not only improve recognition of functional regulatory variants underlying a trait or disease but help us to design better MPRA experiments. The most straightforward tactic would be to empirically exclude tags with a large predicted sequence effect; alternatively, one can theoretically screen large tag sequence libraries to identify sequences we deem to have a significant impact and eliminate them from consideration. As base counts and RBP binding sites are predicted to affect the tag expression significantly, one can design tags with similar profiles of base counts and avoid potential RBP binding sites to systematically reduce tag sequence-specific effects. It is important to note that these binding sites should be considered with the flanking sequences. These strategies may not be feasible for a completely new MPRA design as the tag sequence effect is specific to experiment designs and cell types. Also, it is impossible to apply these strategies to MPRA that uses degenerate tags (Tewhey et al. 2016; Klein et al. 2020). Even in these situations, MTSA can mitigate the problem.

We show here that post-transcriptional mechanisms can affect MPRA expression through tag sequences. However, the sequence features we evaluated were only able to explain <50% of the variation, indicating that additional mechanisms affecting reporter expression may exist. For example, RNA modifications in the 3’UTR, such as adenosine-to-inosine (A-to-I) editing and N^6^-methyladenosine (m^6^A), have been shown to affect mRNA expression (Nishikura 2010; Fu et al. 2014). It is also possible that we may have underestimated the true effects of the evaluated features since there are several assumptions and limitations in our analyses. For example, RBPs not in the CISBP-RNA database or yet unknown RBPs can affect the tag expression. The estimated stability may not capture the full complexity of the *in vivo* contexts, either.

Given that post-transcriptional mechanisms can significantly affect MPRA expression, we may have to be more cautious in interpreting STARR-seq results. Since CRE sequences by themselves are transcribed in STARR-seq (Arnold et al. 2013), they can subsequently affect their expression through post-transcriptional mechanisms in a CRE sequence-specific fashion. We analyzed two STARR-seq data sets for which tags were added in addition to CREs and found significant tag sequence effects similar to the other data sets. However, many tags showed more pronounced effects on its expression, beyond what could be predicted by tag sequences. Specifically, many tags have much lower expression than their predicted expression (**Supplemental Fig. S3A and B**). We speculate that this could result from transcribed CREs, but more careful evaluations are required. Ultimately, a sequence-based understanding of tag effects on reporter expression will help us to better interpret MPRA studies.

Of note, the SVR with gkm-kernel is a generalizable approach and can be applied to other problems, such as quantitative prediction of ChIP-seq, ATAC-seq, and DNase-seq signals from DNA sequences. Thus, we also provide a general implementation of SVR with gkm-kernel in the LSGKM software package.

## METHODS

### Public MPRA data sets

We downloaded public MPRA data sets from the Gene Expression Omnibus (GEO) database for the following studies: GSE31982 (Melnikov et al. 2012), GSE33367 (Kheradpour et al. 2013), GSE87711 (Ulirsch et al. 2016), GSE71279 (Ernst et al. 2016), GSE83894 (Inoue et al. 2017), and GSE142696 (Klein et al. 2020). From the Melnikov study (Mel12), we used the ‘single-hit-scanning’ data set for which multiple tags per CRE were available. From the Kheradpour study, we analyzed both HepG2 (Khe13) and K562 (Khe13K) data sets. From the Ulirsch study, we primarily used the K562 data set (Uli16) for all analyses, but we also analyzed the K562+GATA1 data set (Uli16G) for the functional variant analysis and the SVR weight comparison. From the Ernst study, we were only able to analyze the pilot design MPRA data sets since the tiling design has only one tag per element. We used both HepG2 (Ern16) and K562 (Ern16K) data sets. From the Inoue study, we primarily used the episomal data set (Ino17), but we also analyzed the chromosomal data set (Ino17W) for the SVR weight comparison. From the Klein study, we evaluated the nine MPRA data sets but primarily analyzed the STARR-seq design data sets (HSS and ORI). In addition, we extracted and analyzed the tag-level expression from supplemental data for the CRE-seq: Mog13 (Mogno et al. 2013) and Kwa14 (Kwasnieski et al. 2014). Regarding the Tewhey study (Tewhey et al. 2016), we obtained the tag-level expression data from the author and used the 7.5k design MPRA data sets, NA12878 (Tew16) and NA19239 (Tew16N), as the other one was too sparse at the individual tag level.

For each data set, we first combined all replicates to increase sequence coverage per tag, excluding tags with zero DNA or RNA counts. Expression was calculated as log_2_(RNA/DNA) followed by normalization to zero mean for each of the CREs to obtain relative expression. Next, we excluded tags with small DNA counts to reduce non-sequence-specific variation. The threshold was manually determined for each data set based on the relationship between relative expression and DNA (plasmid) tag counts. For the nine data sets from Klein’s study, we selected tags that have enough number of DNA reads (counts per million reads (CPM)>3) in all nine MPRA data sets for a fair comparison. We further excluded CREs with a small number of tags to reduce the variance of relative expression caused by a small sample size. In total, for training, we identified at least 10,000 tags for most of the data sets. The detailed parameters used for MTSA analyses are summarized in **Supplemental Table S7**.

### Building sequence-based models using Support Vector Regression

To model tag expression from the sequence we used support vector regression (SVR) methods (Drucker et al. 1997) using gapped *k*-mer frequencies from tags and their flanking sequences (±5bp) as features. We used 8-mers with 4 gaps (wildcards) as a default parameter, throughout the study. To implement SVR, we adopted and modified the LS-GKM (Lee 2016) and LibSVM (Chang and Lin 2011) software. To remove CRE-level bias, we first trained an SVR model with 5-fold cross-validation and corrected expression using the trained model. We then retrained the SVR model using corrected expression. We estimated expression from sequence using 5-fold cross-validation and evaluated model performance using Pearson correlation coefficients between observed and predicted relative expression values. We also separately trained an SVR model using all data to score 8-mer weights as well as predict sequence-specific biases of tags excluded from training.

### Analysis of raw tag expression comparisons

Our sequence-based correction was evaluated by comparing individual tag expression within a CRE. Specifically, we randomly selected two tags from each CRE and built the SVR model by following the process described above. We then predicted the sequence effects of the reserved tags using the trained model and corrected raw expression. For a fair comparison, we only kept tag pairs that have the same number of DNA reads (**Supplemental Table S7**).

### Analysis of regulatory variants affecting red blood cell traits

To evaluate the efficacy of our sequence-based correction on regulatory variant identification, we analyzed the MPRA data (Uli16) designed for identifying functional regulatory variants affecting red blood cell traits (Ulirsch et al. 2016). We obtained the original R scripts from the supplemental website of the authors (https://www.bloodgenes.org/RBC_MPRA) and modified them in order to add our MTSA correction routine. We performed analysis with and without MTSA corrections, and identified 37 and 61 variants that showed significantly different CRE activities between reference and alternative alleles, respectively. For the deltaSVM comparison analysis, we obtained pre-calculated deltaSVM scores for all variants available from the supplemental website (Ulirsch et al. 2016). deltaSVM was calculated using a gkm-SVM model trained on K562 DNaseI hypersensitive sites, data that are completely independent of MPRA data. Note that the training sequences for this deltaSVM model are entirely different from the training sequences for the tag correction.

### Analysis of potential molecular mechanisms of tag sequence effects on reporter expression

We considered five different biological feature sets by which tag sequences could affect reporter expression in MPRA. Specifically, we evaluated (1) base count effects on expression (2) tags as CREs themselves, (3) tag effects on the stability of mRNA secondary structures, (4) miRNA binding sites, and (5) RNA-binding protein (RBP) binding sites. The detailed steps to calculate each set of features is described in the following sections. For each feature set, we built a multivariate linear regression model with the predicted tag effects on reporter expression as a dependent variable with and without adjusting the effects of other features and estimated the variance attributed to these features. We also built regression models using all features simultaneously (*i.e.*, a full model) and the two most significant feature sets (RBP + base counts). We used adjusted r^2^ from the lm() function implemented in R software.

### Evaluation of tag sequences as CREs

To quantify CRE activities of tags, we computationally predicted tag sequences’ CRE activities using gkm-SVM models trained on DNase-seq data from the same cell-type. We followed our previously established pipeline with minor changes (Lee et al. 2018). Briefly, starting from the top 50,000 regions from the ENCODE DNase-seq narrowPeak files (ENCFF711IED for HepG2, ENCFF821KDJ for K562, ENCFF127KSH for HEK293T, and ENCFF073ORT for GM12878), we trained gkm-SVM models against equal numbers of random genomic regions that match GC content and repeat fraction of the positive set. We used 600bp regions extended from the centers of peaks. All three models achieved high accuracy, as measured by area under the ROC curves (AUC: 0.92-0.94) with 5-fold cross-validation. We then scored tags with ±5bp flanking sequences using the model from the same cell-type (HEK293T for Mel12, HepG2 for Khe13/Ino17, and K562 for Uli16).

### Minimum free energy calculation of mRNA secondary structure

To evaluate the effect of tag sequences on mRNA stability of the secondary structure, we calculated minimum free energy (MFE) of tag sequences with ±50bp flanking sequences. We used the “RNAfold” command-line tool implemented in the Vienna RNA software (Gruber et al. 2008).

### Analysis of miRNA binding sites in tag sequences

To identify functional miRNA binding sites in the tag sequences, we first determined the top 20 highly expressed miRNA families as defined by those sharing the same miRNA 7-bp seed in each of the four cell types (HEK293T, HepG2, K562, and a lymphoblastoid cell line [GM12878]), since miRNA expression is mainly cell-type specific. We obtained miRNA expression data for K562, HepG2, and GM12878 from ENCODE (accession no: ENCSR569QVM for K562 and ENCSR730NEO for HepG2, ENCSR770HBF for GM12878), and HEK293T from miRmine (accession no: SRX556516) (Panwar et al. 2017). For each miRNA in the corresponding cell type, we identified tags with ±5bp flanking sequences containing its 7bp miRNA seed, resulting in a binary variable. We obtained miRNA seed sequences from TargetScan 7.2 (Agarwal et al. 2015). Rare miRNA binding sites (<5 matches of tags) were removed. To remove redundant miRNAs, we calculated the pairwise Pearson correlation and only kept miRNAs with r <0.8. The number of miRNA variables selected for each data set is summarized in **Supplemental Table S8**.

### Analysis of binding sites of RNA-binding proteins (RBP) in tag sequences

To identify putative binding sites of RNA-binding proteins (RBP) within the tag sequences, we analyzed 172 motifs associated with 143 distinct RBPs from the CISBP-RNA database (Ray et al. 2013). For each of these motifs, we scanned tags with ±5bp flanking sequences using FIMO (Grant et al. 2011) with the P < 0.001. Similar to miRNA, we only included highly expressed RBPs (TPM>10) and removed redundant motifs with cutoff r <0.8. Rare RPB binding sites (<1% matches of tags) were also removed. The number of RBP variables selected for each data set is summarized in **Supplemental Table S8**.

## Availability and implementation

We implemented new software, <u>M</u>PRA <u>T</u>ag <u>S</u>equence <u>A</u>nalysis (MTSA), which is freely available under the GNU General Public License v3.0 as Supplemental Material and at the GitHub repository: https://github.com/Dongwon-Lee/mtsa.

A generic implementation of SVR with gkm-kernel is also freely available at the LSGKM GitHub repository: http://github.com/Dongwon-Lee/lsgkm.

## ACKNOWLEDGMENTS

We thank Drs. Vijay Sankaran and Jacob Ulirsch for discussions and assistance in analyzing their published MPRA data set. We thank Dr. Ryan Tewhey for kindly providing us the tag-level expression data. This study has benefited from constructive comments from members of the Chakravarti laboratory.

## FUNDING

The research reported here was supported by the computational resources of the High-Performance Cluster (BigPurple) at NYU School of Medicine and NIH grants GM104469, HL086694, and HL128782.

## AUTHOR CONTRIBUTIONS

D.L. and A.K. conceived the study; D.L. developed the method and performed all the analyses; A.K., C.L., M.M., M.A.B., and A.C. contributed to methods development; A.K., C.L., M.A.B., and A.C. contributed to data analysis; D.L. and A.C wrote the manuscript; all authors were involved in manuscript revision.

## COMPETING FINANCIAL INTERESTS

The authors declare no competing financial interests.

## Notes

### Competing Interest Statement

The authors have declared no competing interest.

