## Supplemental Material for "Sequence-based correction of barcode bias in massively parallel reporter assays"

**This document includes:**

Supplemental Figures S1 to S10

Supplemental Tables S1 to S7

Supplemental Notes

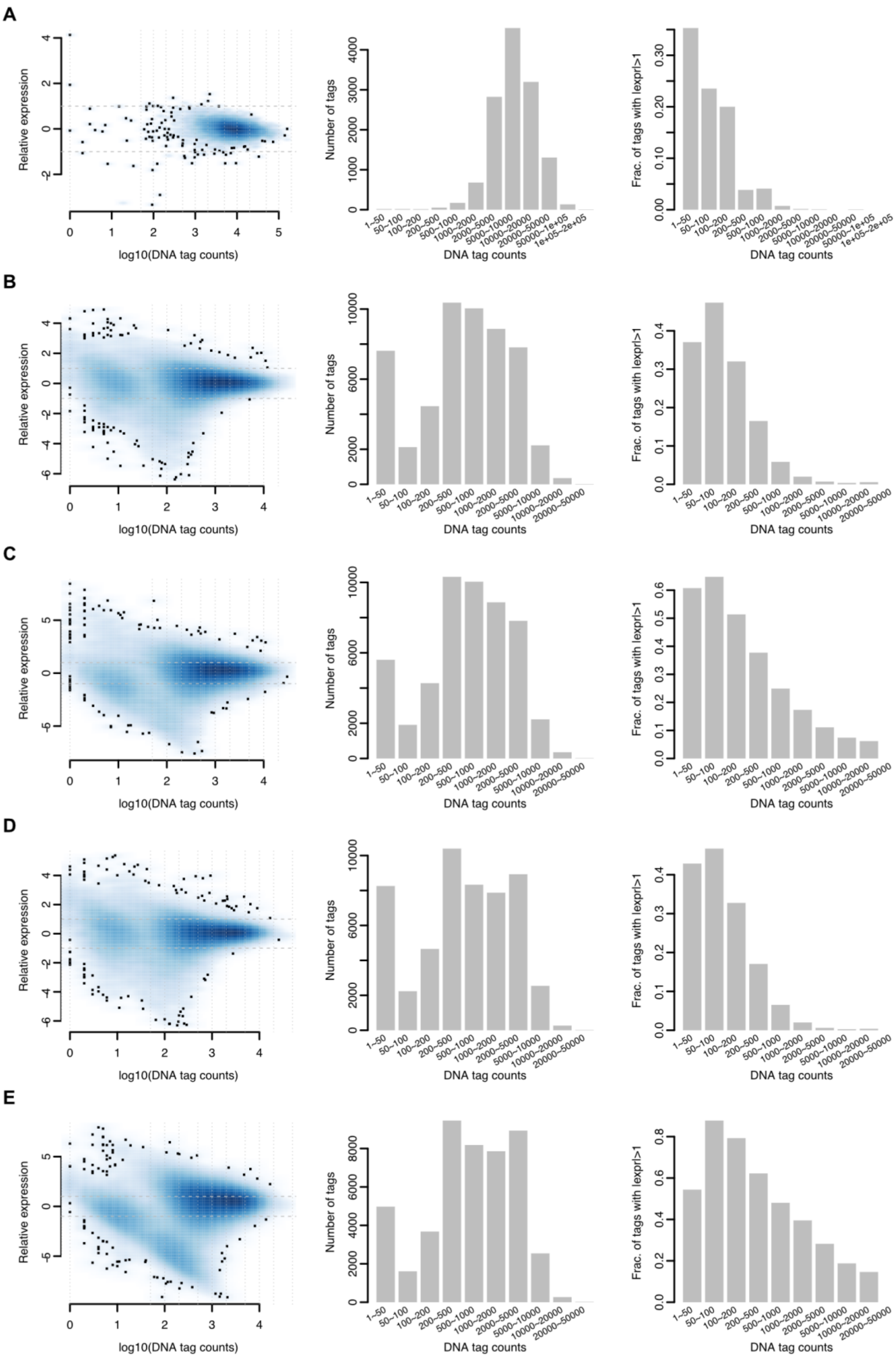

A
Relative expression
log10(DNA tag counts)
Number of tags
DNA tag counts
Frac. of tags with |expr|>1
1~50
50~100
100~200
200~500
500~1000
1000~2000
2000~5000
5000~10000
10000~20000
20000~50000
50000~1e+05
1e+05~2e+05
B
Relative expression
log10(DNA tag counts)
Number of tags
DNA tag counts
Frac. of tags with |expr|>1
C
Relative expression
log10(DNA tag counts)
Number of tags
DNA tag counts
Frac. of tags with |expr|>1
D
Relative expression
log10(DNA tag counts)
Number of tags
DNA tag counts
Frac. of tags with |expr|>1
E
Relative expression
log10(DNA tag counts)
Number of tags
DNA tag counts
Frac. of tags with |expr|>1

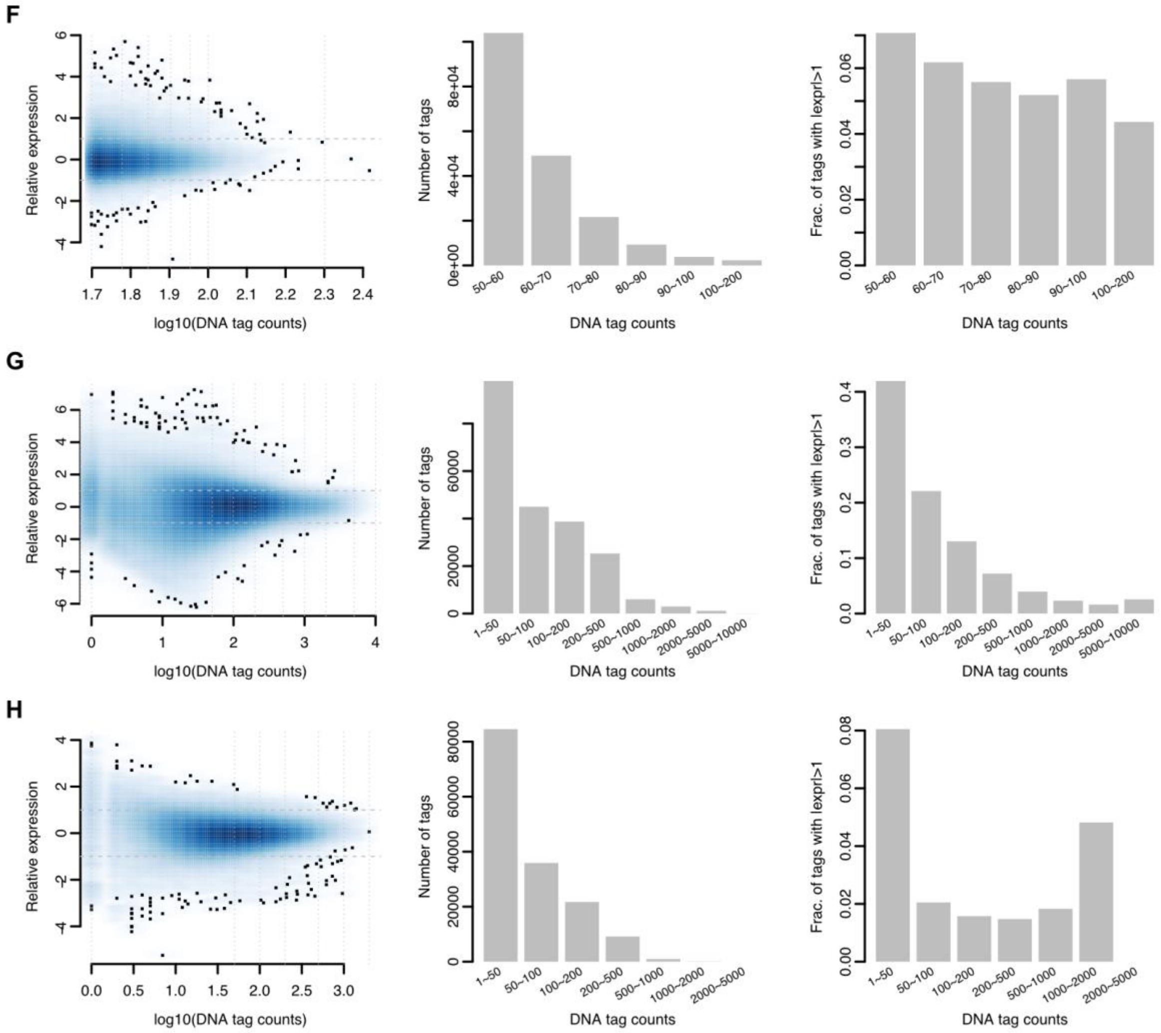

**Supplemental Figure S1: DNA tags with low read counts are more variable.** The relationships between DNA (plasmid) read counts and their relative expression **(1st column)**, tag frequencies binned by their DNA read counts **(2nd column)**, and tag fractions with at least two-fold relative expression binned by their DNA read counts **(3rd column)** from the Melnikov *et al.* 2012 **(A, Mel12)**, Kheradpour *et al.* 2013 for HepG2 **(B, Khe13)** and K562 **(C, Khe13K)**, Ernst *et al.* 2016 for HepG2 **(D, Ern16)** and K562 **(E, Ern16K)**, Tewhey *et al.* 2016 **(F, Tew16)**, Ulirsch *et al.* 2016 **(G, Uli16)**, and Inoue *et al.* 2017 **(H, Ino17)** studies.

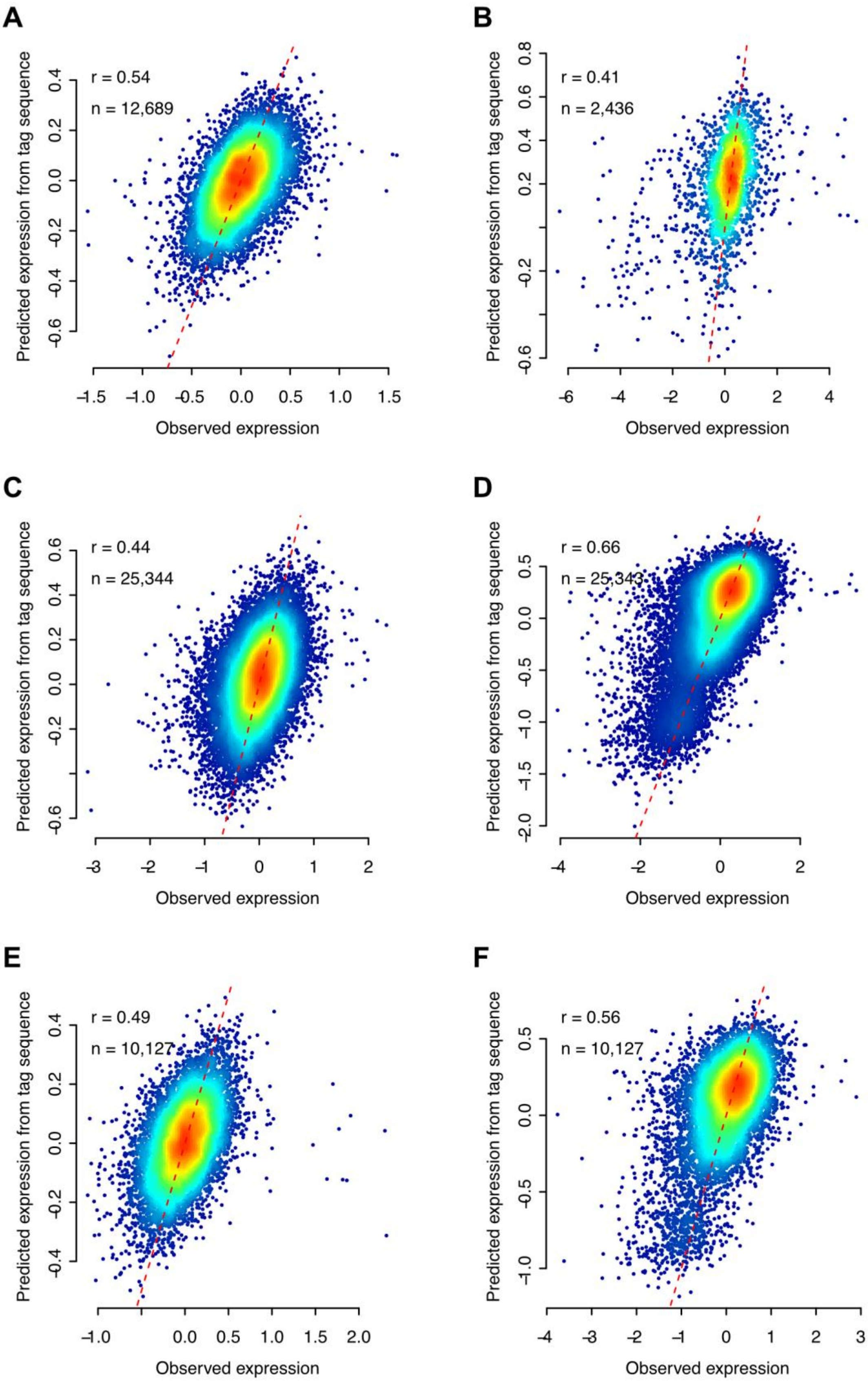
A
r = 0.54
n = 12,689
B
r = 0.41
n = 2,436
C
r = 0.44
n = 25,344
D
r = 0.66
n = 25,343
E
r = 0.49
n = 10,127
F
r = 0.56
n = 10,127
Predicted expression from tag sequence
Observed expression

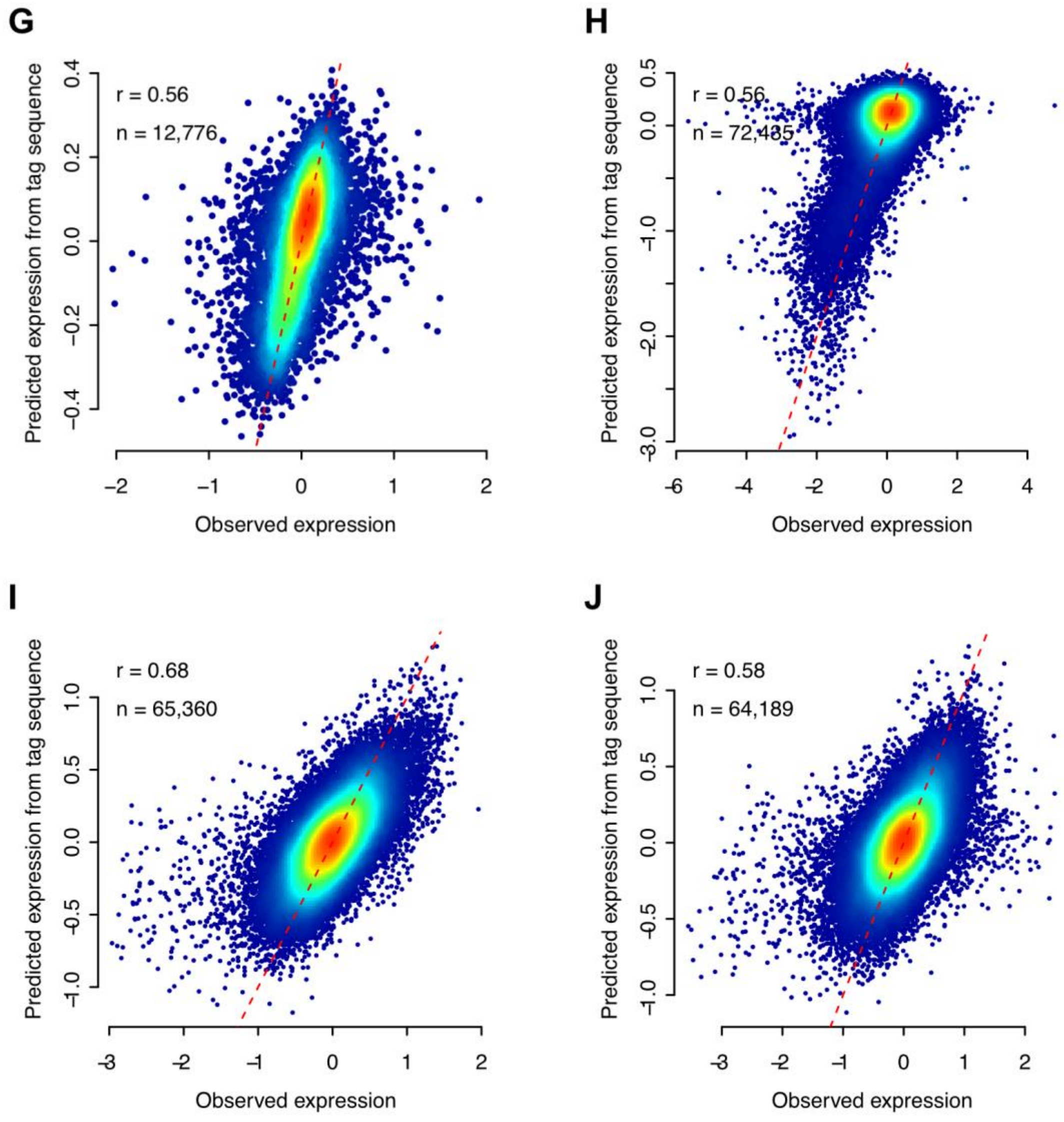

**Supplemental Figure S2: Support vector regression accurately predicts tag sequence effects.** The observed relative expression (X-axis) is compared to the predicted expression from tag sequence (Y-axis) using 5-fold cross-validation for the data in Melnikov *et al.* 2012 **(A, Mel12)**, Mogno *et al.* 2013 **(B, Mog13)** Kheradpour *et al.* 2013 for HepG2 **(C, Khe13)**, and K562 **(D, Khe13K)**, Ernst *et al.* 2016 for HepG2 **(E, Ern16)** and K562 **(F, Ern16K)**, Kwasnieski et al. 2014, **(G, Kwa14)** Tewhey *et al.* 2016 **(H, Tew16)** and Inoue *et al.* 2017 using episomal **(I, Ino17)** and integrated **(J, Ino17W)** constructs, respectively. Pearson correlations (*r*) and numbers of data points (*n*) are shown. Red dashed lines indicate the Y = X line. The density of data points is represented by a color heatmap calculated by a bivariate Gaussian Kernel Density Estimate function with default parameters (see: densCols() and bkde2D() functions in R for details).

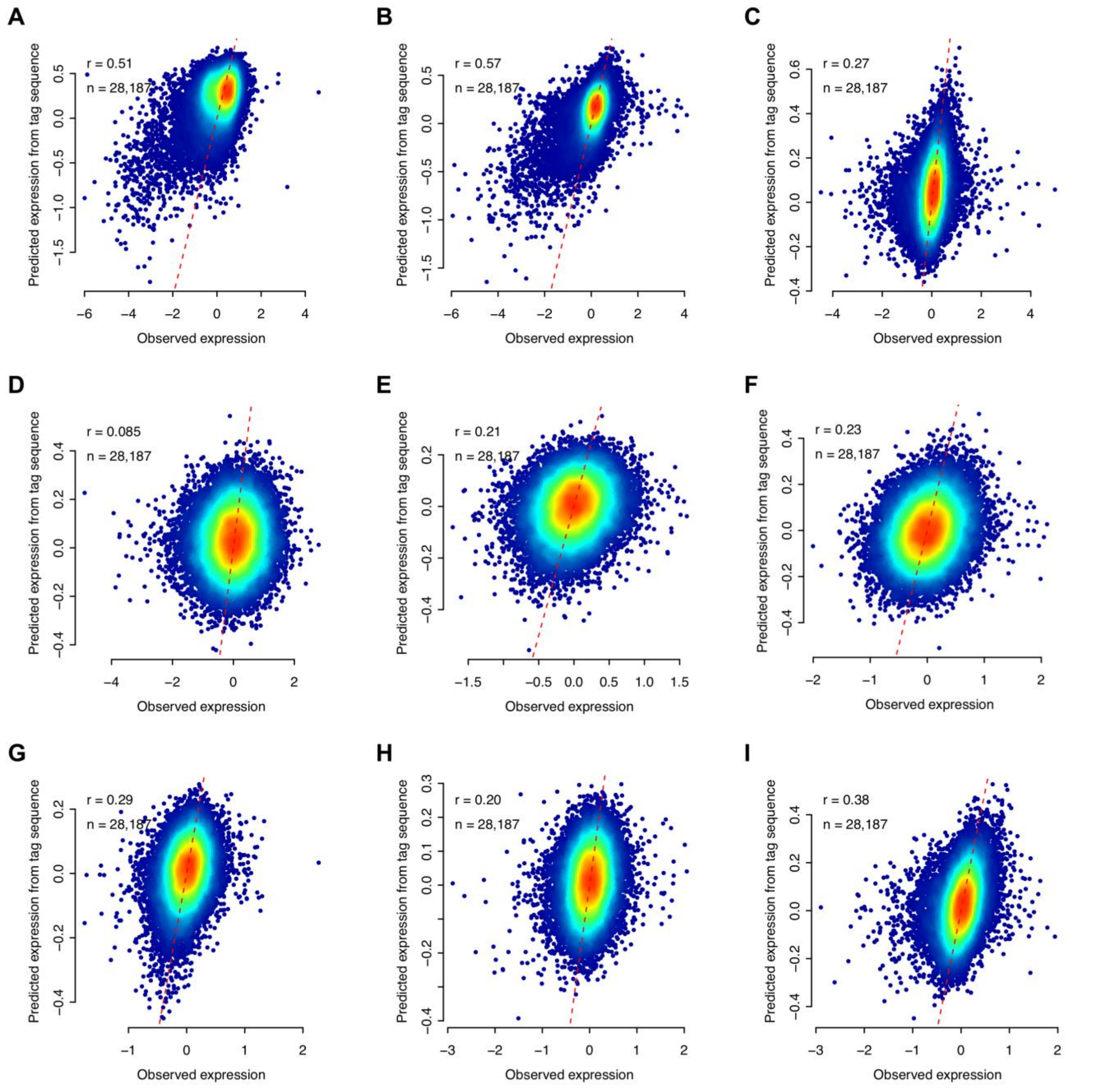

**Supplemental Figure S3: MTSA analysis of the nine MPRA data sets from Klein et al. 2020.** Observed relative expression are compared to MTSA predicted expression from 5-fold cross-validation for the original STARR-seq **(A, HSS)**; STARR-seq with no minimal promoter **(B, ORI)**; pGL4.23c vector **(C, pGL4)**; lentiMPRAs with both the enhancer library and barcodes in the 3' UTR of the reporter packaged with MT integrase **(D, 3'/3' MT)** and WT integrase **(E, 3'/3' WT)**; lentiMPRAs with the enhancer library upstream of the minimal promoter and the associated barcodes in the 3′ UTR of the reporter gene with MT integrase **(F, 5'/3' MT)** and WT integrase **(G, 5'/3' WT)**; lentiMPRAs with the enhancer library upstream of the minimal promoter and barcodes in the 5' UTR of the reporter with MT integrase **(H, 5'/5' MT)** and WT integrase **(I, 5'/5' WT)**.

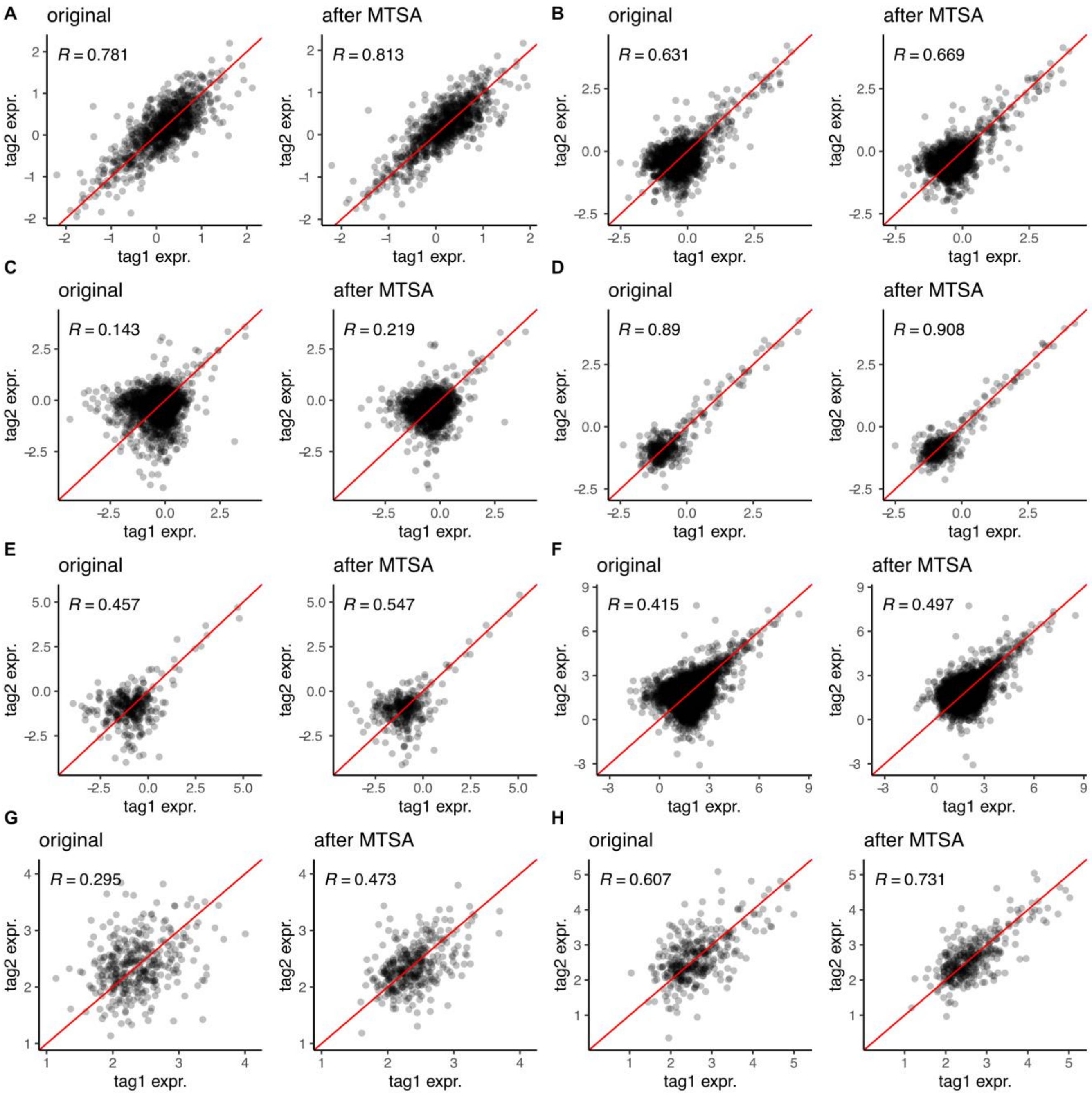

**Supplemental Figure S4: Tag sequence-based correction reduces variation within CREs.** Scatter plots of tag expression of two randomly selected tags within CREs are shown with and without MTSA correction in a $\log_2$ scale, for the data in Mel12 **(A)**, Khe13 **(B)** and Khe13K **(C)**, Ern16 **(D),** Ern16K **(E)**, Tew16 **(F),** Ino17 **(G),** and Ino17W **(H)**, respectively. *R* is the Pearson correlation coefficient, and the red dashed line indicates the Y = X line.

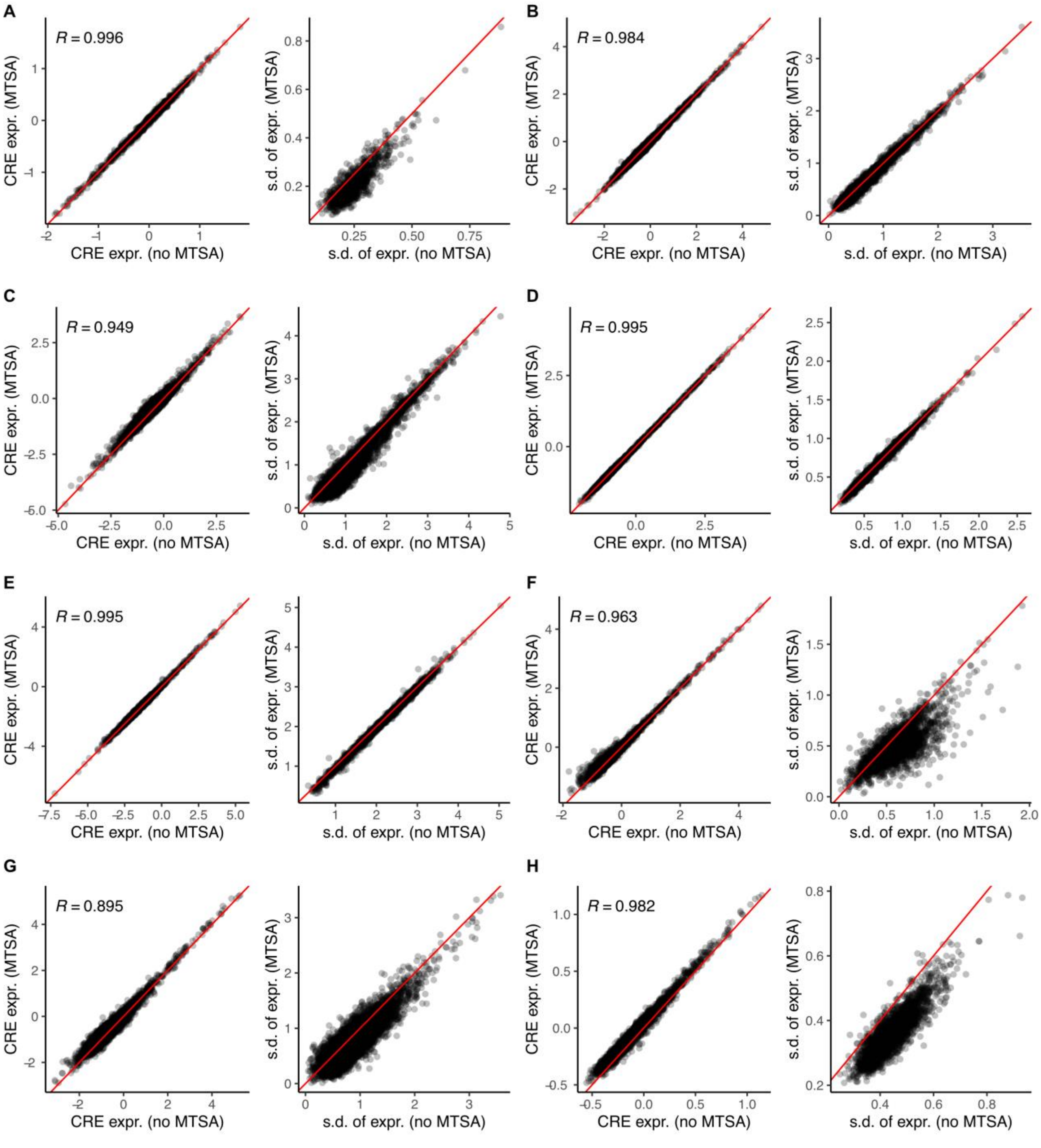

**Supplemental Figure S5: MTSA correction marginally affects CRE expression but significantly reduces its variance across tags.** The correlation between CRE-level expression with and without MTSA correction **(the 1st and 3rd column)** and their standard deviation **(the 2nd and 4th column)** are shown for the data in Mel12 **(A)**, Khe13 **(B)**, Khe13K **(C)**, Ern16 **(D)**, Ern16K **(E)**, Tew16 **(F),** Uli16 **(G)**, and Ino17 **(H)**. *R* is the Spearman's correlation, and the red line indicates the Y = X line. The average of $\log_2$(RNA/DNA) across tags is the CRE-level expression. We only used CREs with at least 3 tags and tags with at least one read per million.

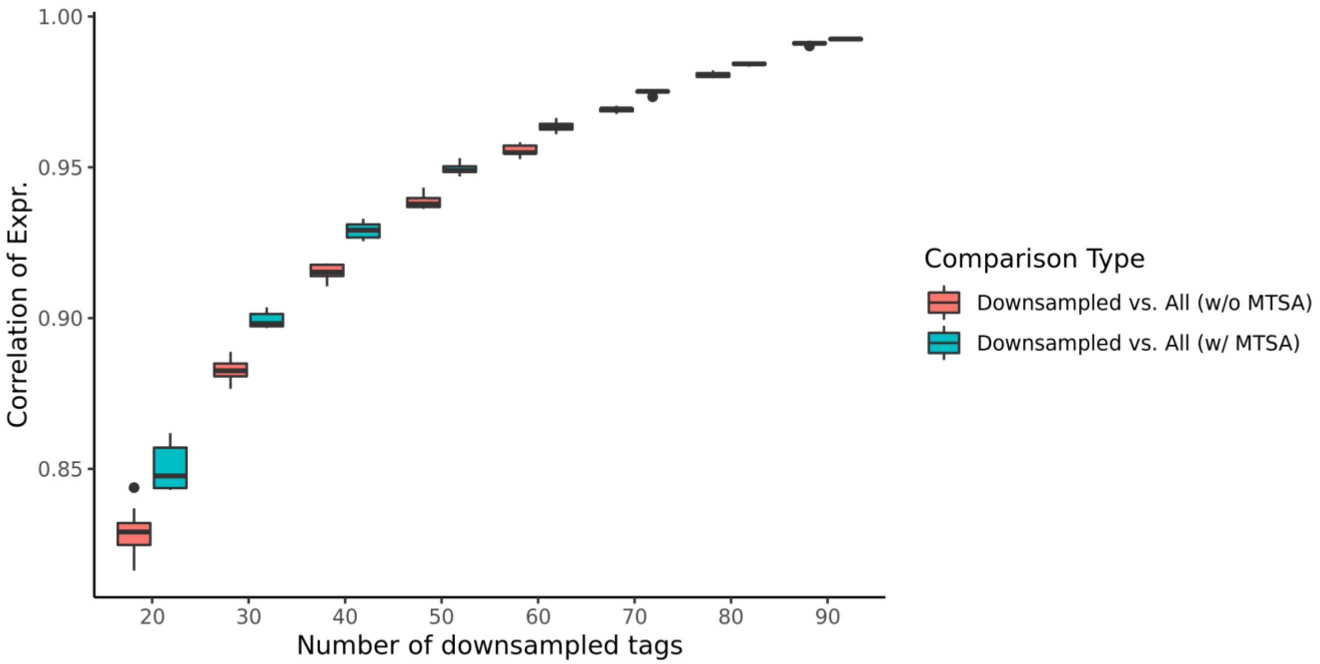

**Supplemental Figure S6: Correlation of expression between down-sampled tags and all tags before and after MTSA correction.** For each of the down-sampled tags with n=20, 30, ..., 90, we calculated CRE-level expression and compared them to those calculated using all tags (n=100) with and without MTSA correction. MTSA correction was applied to both. For each tag number, we repeated 10 times and calculated average Spearman correlation coefficients. Improvement achieved by MTSA correction is gradually attenuated and the average improvement is <2% when n>50. The average of $\log_2$(RNA/DNA) across tags is the CRE-level expression. We only used CREs with at least three tags, and tags with at least one read per million.

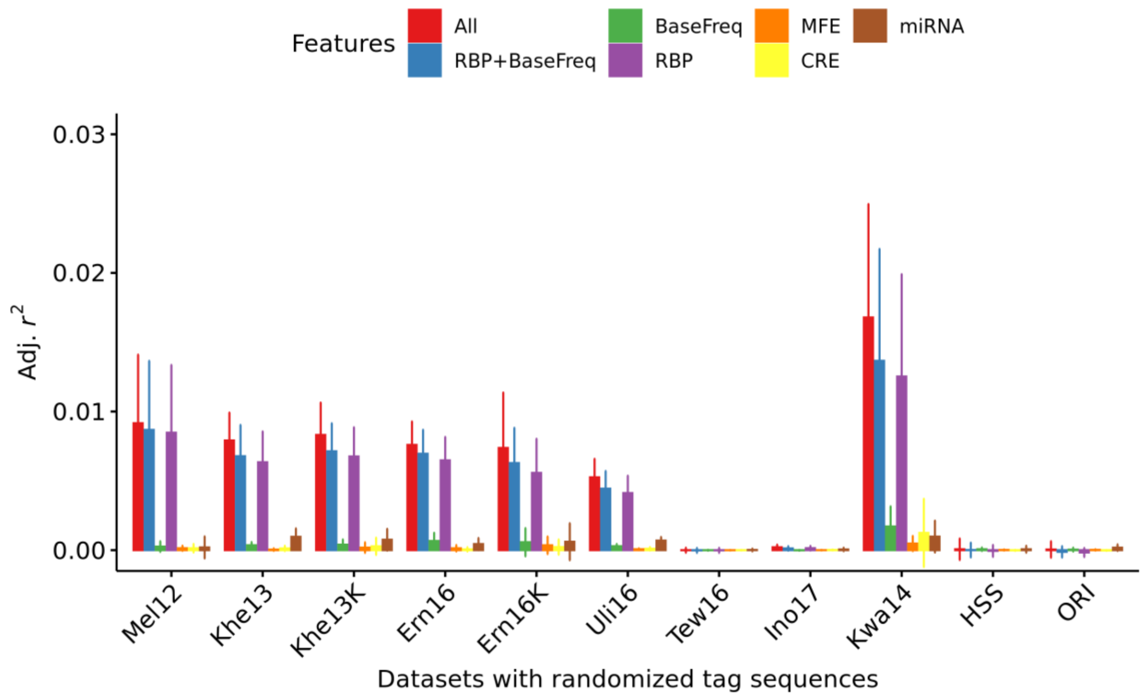

**Supplemental Figure S7: Randomized tag sequences do not explain the variation of relative tag expression attributed to specific biological features.** As a negative control, we randomized the tag sequences and performed the same multivariate linear regression analysis as **Fig. 5**. We repeated the analysis ten times to estimate the mean and the standard deviation of adjusted $r^2$. We found that <1% of the variance can be explained by the randomized tags for all data sets, except Kwa14.

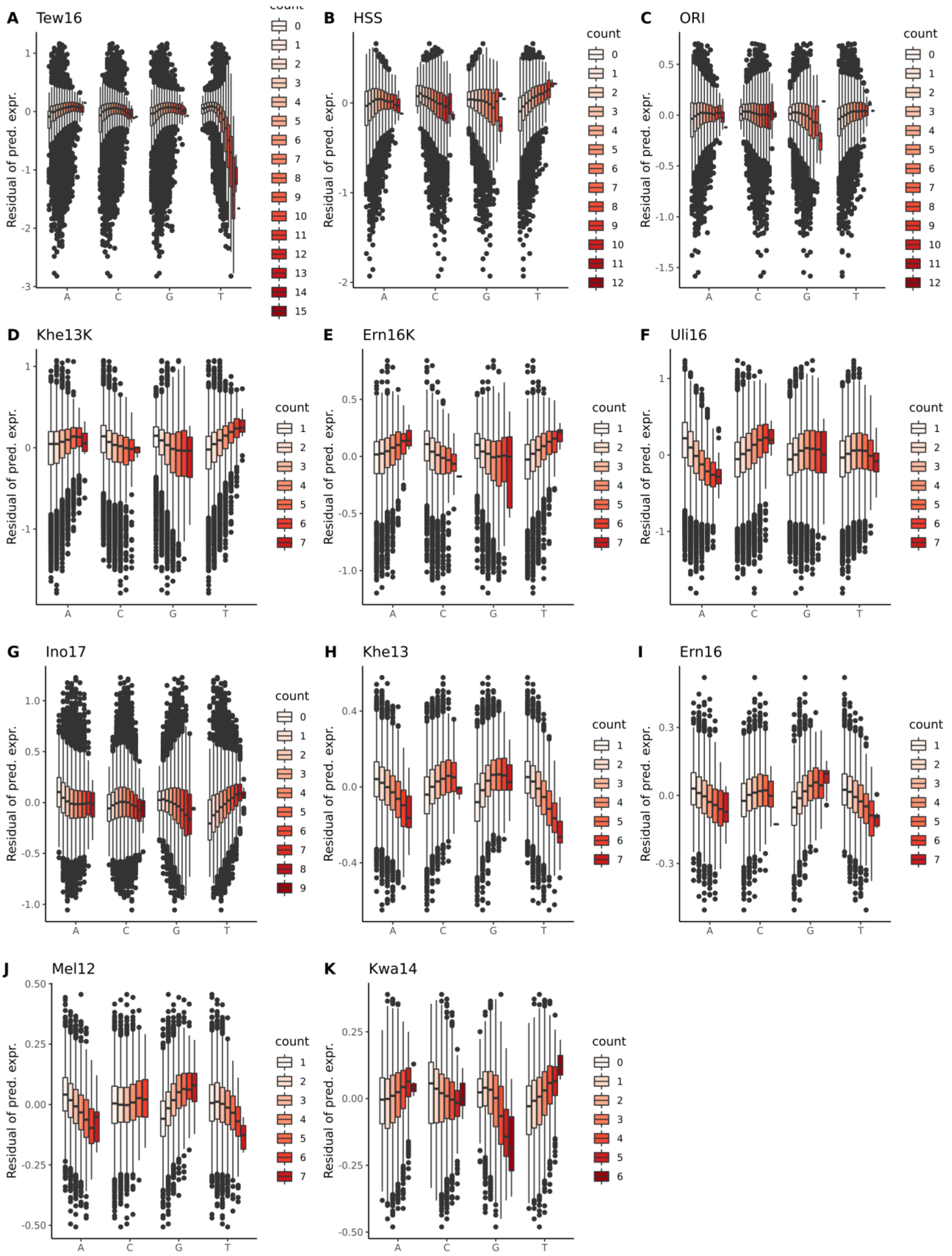
A Tew16
B HSS
C ORI
D Khe13K
E Ern16K
F Uli16
G Ino17
H Khe13
I Ern16
J Mel12
K Kwa14
Residual of pred. expr.
count
A C G T

**Supplemental Figure S8: Effect of base counts on relative expression is specific to experimental designs and cell types.** Distributions of residuals of the predicted relative expression are stratified by base counts for Tew16 **(A)**, HSS **(B)**, and ORI **(C)**, Khe13K **(D)**, Ern16K **(E)**, Uli16 **(F)**, Ino17 **(G)**, Khe13 **(H)**, Ern16 **(I)**, Mel12 **(J)**, Kwa14 **(K)**. In general, the count of T shows the strongest effect on the predicted expression, although the direction of effect is specific to the experiment and cell types. Data sets associated with long tags (especially created by degenerate bases) or measured in K562 **(A-G)** tend to have more extreme outliers than other data sets **(H-K)**.

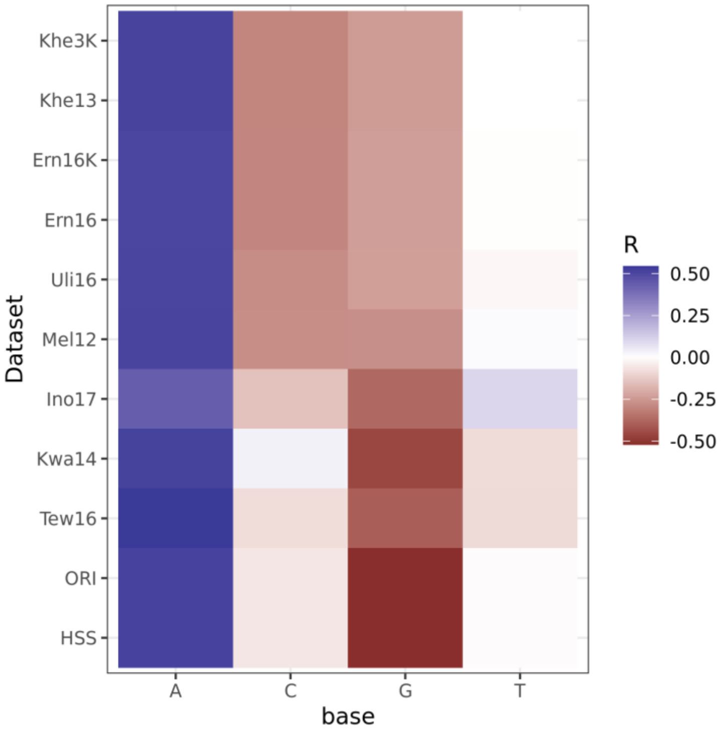

**Supplemental Figure S9: Base counts in tags strongly correlate with minimum free energy of mRNA secondary structure.** For each of the data sets, correlations of MFE and each of the base counts in tags are shown as a heatmap. Expectedly, MFE is strongly positively correlated with the count of 'A' for all data sets, while negatively correlated with 'C', and 'G' counts.

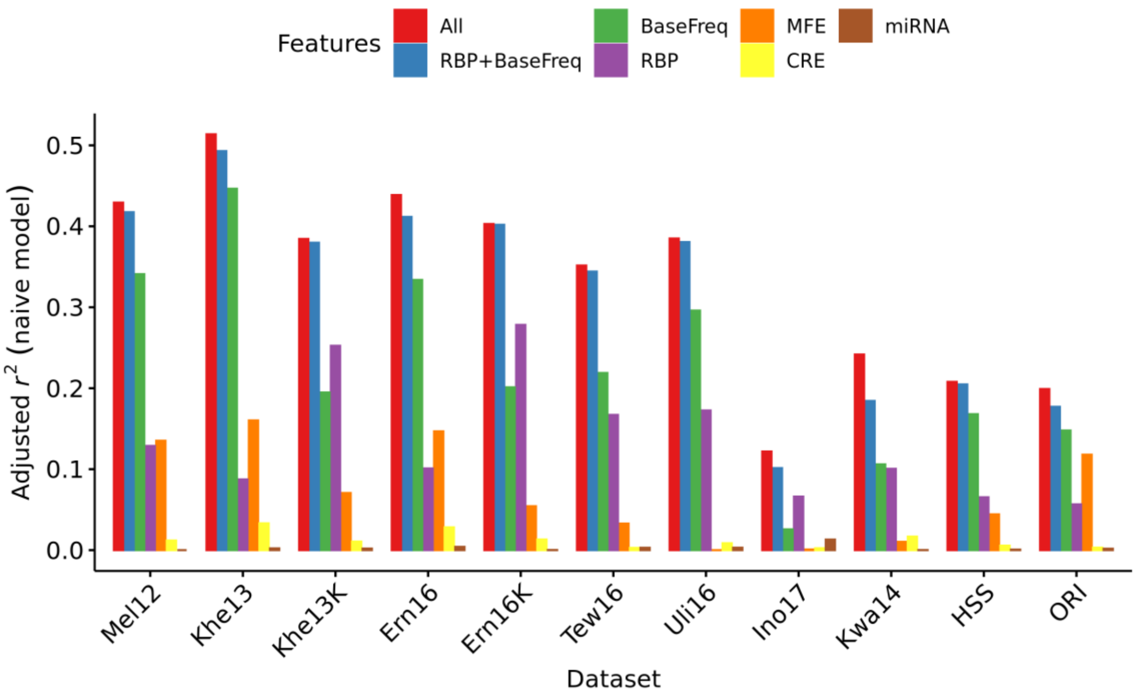

**Supplemental Figure S10: Base counts capture signals that can be explained by other potential biological mechanisms.** Multivariate linear regression analyses were performed similar to **Fig. 5**, except that the dependent variable (predicted relative expression) was not adjusted by other features sets. The BaseFreq feature set captures more variation in these naïve models than the residual models in **Fig. 5** because it captures variations explained by other biological features (i.e., MFE, RBP, and CRE). Similarly, MFE explains up to 15% of variation in expression if it is not adjusted by BaseFreq. **BaseFreq**: base frequencies in tag sequences, **MFE**: minimum free energy of mRNA secondary structures, **CRE**: tags as cis-regulatory elements, **miRNA**: microRNA binding sites, **RBP**: RNA binding protein binding sites, **RBP+BaseFreq**: RBP and BaseFreq features combined, **All**: all features combined.

| Data set ID | Year | Cell Type | # of CREs | Tags per CRE | # of Constructs | Oligo Length | CRE Length | Tag Length | Restriction Sites | Reporter Gene |
|---|---|---|---|---|---|---|---|---|---|---|
| **Mel12** | 2012 | HEK293 | 1,000 | 13 | 13,000 | 142 | 87 | 10 | KpnI – XbaI | Luciferase |
| **Khe13** | 2013 | HepG2, K562 | 5,400 | 10 | 54,000 | 200 | 145 | 10 | KpnI – XbaI | Luciferase |
| **Mog13** | 2013 | Yeast (SC) | 2,467 | Var. | 6,908 | . | Var. | 15 | EagI – XbaI | YFP |
| **Kwa14** | 2014 | K562 | 3,237 | 4 | 12,931 | 200 | 130 | 9 | XhoI – SphI | DsRed |
| **Ern16** | 2016 | HepG2, K562 | 2,250 | 24 | 54,000 | 200 | 145 | 10 | KpnI – XbaI | Luciferase |
| **Tew16** | 2016 | LCLs | 7,500 | Var. | ~3.4M | 180 | 150 | 20 | . | GFP |
| **Uli16** | 2016 | K562 | 16,534 | 14 | 231,476 | 200 | 145 | 10 | KpnI – XbaI | Luciferase |
| **Ino17** | 2017 | HepG2 | 2,440 | 100 | 244,000 | 230 | 171 | 15 | SbfI – EcoRI | EGFP |
| **Kle20** | 2020 | HepG2 | 2,440 | Var. | 4.1M ~ 9.8M | 230 | 171 | 15 | SbfI – EcoRI | GFP |

**Supplemental Table S1: Design details of the public MPRA data sets evaluated in this study.** Details of nine different MPRA studies are summarized. Studies that used variable number of tags per CRE are marked as "Var." SC: Saccharomyces cerevisiae; YFP: yellow fluorescent protein; DsRed: red fluorescent protein; EGFP: enhanced green fluorescent protein; LCLs; lymphoblastoid cell lines.

| Data set ID | Correlation after 1st training | Correlation after 2nd training | Difference in Correlation |
|---|---|---|---|
| **Mel12** | 0.50 | 0.54 | +0.04 |
| **Mog13** | 0.39 | 0.41 | +0.02 |
| **Khe13** | 0.38 | 0.44 | +0.06 |
| **Khe13K** | 0.59 | 0.66 | +0.07 |
| **Kwa14** | 0.44 | 0.56 | +0.12 |
| **Ern16** | 0.44 | 0.49 | +0.05 |
| **Ern16K** | 0.51 | 0.56 | +0.05 |
| **Tew16** | 0.54 | 0.56 | +0.02 |
| **Tew16N** | 0.28 | 0.30 | +0.02 |
| **Uli16** | 0.65 | 0.71 | +0.06 |
| **Ino17** | 0.66 | 0.68 | +0.02 |
| **Ino17W** | 0.57 | 0.58 | +0.01 |

**Supplemental Table S2: Model performance is consistently improved by bias correction based on the initial training.** For each of the data sets, we calculated correlations between the observed relative expression and the predicted expression after the first training (2nd column) as well as the second training (3rd column). The correlation values in the 3rd column are the same as the *r* values shown in **Fig. 2A** and **Supplemental Fig. S2**. Differences in the correlations are shown in the 4th column.

| Data set ID | Correlation using reverse complement sequences as the same features | Correlation using reverse complement sequences as distinct features | Difference in Correlation |
|---|---|---|---|
| **Mel12** | 0.53 | 0.54 | +0.01 |
| **Mog13** | 0.22 | 0.41 | +0.19 |
| **Khe13** | 0.43 | 0.44 | +0.01 |
| **Khe 13K** | 0.65 | 0.66 | +0.01 |
| **Kwa14** | 0.55 | 0.56 | +0.01 |
| **Ern16** | 0.48 | 0.49 | +0.01 |
| **Ern16K** | 0.55 | 0.56 | +0.01 |
| **Tew16** | 0.42 | 0.56 | +0.14 |
| **Tew16N** | 0.22 | 0.30 | +0.08 |
| **Uli16** | 0.70 | 0.71 | +0.01 |
| **Ino17** | 0.64 | 0.68 | +0.04 |
| **Ino17W** | 0.55 | 0.58 | +0.03 |

**Supplemental Table S3: Using reverse complement sequences as distinct features marginally but consistently improves model performance.** MTSA models were retrained by treating the reverse complement gapped k-mer pairs as the same features, with an assumption that DNA sequences, rather than mRNA sequences, play a major role in affecting the tag expression. The correlation between the observed relative expression and the predicted expression by these models are shown in the 2$^{nd}$ column. The correlation values in the 3$^{rd}$ column are the same as the *r* values shown in **Fig. 2A** and **Supplemental Fig. S2**. Differences in the correlations are shown in the 4$^{th}$ column.

| Data set ID | Correlation of CREs between replicates before MTSA correction | Correlation of CREs between replicates after MTSA correction | Difference in Correlation |
|---|---|---|---|
| **Mel12** | 0.78 | 0.78 | 0 |
| **Khe13** | 0.73 | 0.72 | -0.01 |
| **Khe 13K** | 0.45 | 0.44 | -0.01 |
| **Ern16** | 0.92 | 0.92 | 0 |
| **Ern16K** | 0.70 | 0.70 | 0 |
| **Uli16** | 0.58 | 0.58 | 0 |
| **Ino17** | 0.98 | 0.97 | -0.01 |
| **Ino17W** | 0.99 | 0.99 | 0 |

**Supplemental Table S4: MTSA correction does not improve the correlation of CRE level expression between experimental replicates.** Correlation of CRE-level expression between replicates before and after MTSA correction are shown for the eight different MPRA data sets. The average of $\log_2$(RNA/DNA) across tags was used as the CRE-level expression. We only considered CREs with at least three tags, and tags with at least one read counts per million.

| Mel12 | | | Khe13 | | | Khe13K | | |
|---|---|---|---|---|---|---|---|---|
| **8-mer** | **SVRW** | **RBP** | **8-mer** | **SVRW** | **RBP** | **8-mer** | **SVRW** | **RBP** |
| TCGAGATC | -0.34 | M348_0.6 | ATATATAA | -0.42 | M055_0.6 | TCGAGATC | -2.17 | M348_0.6 |
| AAATATAT | -0.33 | . | ATAAATAA | -0.42 | . | ACGAGATC | -1.82 | M348_0.6 |
| CCACGAGA | -0.32 | . | TAGATATA | -0.4 | . | CCGAGATC | -1.79 | . |
| AAATAAAT | -0.31 | M176_0.6 | TATATATA | -0.4 | M055_0.6 | GCGAGATC | -1.78 | . |
| **CTAGA**AAA | -0.3 | . | AAAAATTA | -0.38 | . | CTCGAGAT | -1.22 | M348_0.6 |
| GAAGATTC | -0.3 | . | ATAAATTA | -0.37 | . | CCCGAGAT | -1.21 | . |
| AAATAATT | -0.3 | . | TATAAATA | -0.37 | . | CGCGAGAT | -1.17 | . |
| AAAAATAT | -0.3 | . | AATAATTA | -0.37 | M001_0.6 | CACGAGAT | -1.16 | M348_0.6 |
| ATATATAT | -0.29 | M055_0.6 | AAAAATAA | -0.37 | . | TTCGAGAT | -1.15 | M348_0.6 |
| ATATAAAT | -0.29 | . | ATAGATAA | -0.36 | M210_0.6 | CCTCGAGA | -1.14 | . |
| AGATGAGA | 0.21 | . | GAGAGGCC | 0.32 | . | GAGATCAA | 0.63 | . |
| AGTGGCCT | 0.21 | . | AAAAGGCC | 0.32 | . | GAAAGATC | 0.63 | M250_0.6 |
| AGAGGTCC | 0.22 | . | AAAATGGC | 0.33 | . | TAAAGATC | 0.7 | M250_0.6 |
| AGTGAGAA | 0.22 | . | AGAGGCCC | 0.35 | . | AAAAGATC | 0.7 | M148_0.6 |
| **CTAGA**GCT | 0.23 | . | AGAGGCCT | 0.35 | . | CGAGATCA | 0.7 | . |
| AGAGGCCG | 0.23 | . | AGAAGCCG | 0.35 | . | AGAGATCA | 0.71 | . |
| **CTAGA**GGC | 0.24 | . | AGAGGCCG | 0.36 | . | AGAGATCT | 0.72 | . |
| TAGAGGCC | 0.26 | . | **CTAGA**GGC | 0.4 | . | GGAAGATC | 0.74 | M250_0.6 |
| AGAGGCCT | 0.26 | . | TAGAGGCC | 0.41 | . | AGAGATCC | 0.78 | . |
| **CTAGA**GCC | 0.34 | . | **CTAGA**GCC | 0.42 | . | AGAGATCG | 0.79 | . |

| Ern16 | | | Ern16K | | | Tew16 | | |
|---|---|---|---|---|---|---|---|---|
| **8-mer** | **SVRW** | **RBP** | **8-mer** | **SVRW** | **RBP** | **8-mer** | **SVRW** | **RBP** |
| ATATATAT | -0.35 | M055_0.6 | TCGAGATC | -1.61 | M348_0.6 | TTTTTTTT | -2.54 | M012_0.6 |
| TATATATA | -0.35 | M055_0.6 | ACGAGATC | -1.43 | M348_0.6 | ATTTTTTT | -2.42 | M012_0.6 |
| ATATATAA | -0.34 | M055_0.6 | GCGAGATC | -1.4 | . | TTTTTTTA | -2.27 | M012_0.6 |
| AAATATAT | -0.33 | . | CCGAGATC | -1.31 | . | AATTTTTT | -2.15 | M025_0.6 |
| ATATATTT | -0.32 | . | GTCGAGAT | -0.94 | M348_0.6 | TATTTTTT | -2.09 | M012_0.6 |
| ATAAATAA | -0.32 | . | TCGAGAGC | -0.87 | . | CTTTTTTT | -2.09 | M012_0.6 |
| AGATATAT | -0.31 | M233_0.6 | TTCGAGAT | -0.87 | M348_0.6 | TTTTTTTG | -2.05 | M012_0.6 |
| ATATATAG | -0.3 | M055_0.6 | ACGAGAGC | -0.85 | . | ACTTTTTT | -1.98 | M012_0.6 |
| AATATATT | -0.3 | . | GGCGAGAT | -0.8 | . | TCTTTTTT | -1.93 | M012_0.6 |
| ATATATTA | -0.3 | . | CCCGAGAT | -0.79 | . | TTTTTTAT | -1.82 | M075_0.6 |
| AGCCGGAA | 0.22 | M290_0.6 | CAAAGATC | 0.44 | M250_0.6 | AGGGGCGG | 0.25 | . |
| AAAAGGCC | 0.23 | . | GGAAAGAT | 0.44 | . | CTAAAATG | 0.25 | M060_0.6 |
| AGAAGCCG | 0.23 | . | GGGAAGAT | 0.45 | . | AAGGGCGG | 0.25 | . |
| AGAGCCCG | 0.25 | . | GAGAAGAT | 0.45 | M148_0.6 | GACGGTGC | 0.25 | . |
| AGAGGCCC | 0.25 | . | TATAGATC | 0.48 | . | ATTTGGAA | 0.25 | M290_0.6 |
| AGAGGCCT | 0.29 | . | AGAGATCC | 0.49 | . | GGGCGGGC | 0.26 | M151_0.6 |
| AGAGGCCG | 0.29 | . | GGAAGATC | 0.52 | M250_0.6 | TGGGCGGA | 0.28 | M065_0.6 |
| TAGAGGCC | 0.33 | . | TAAAGATC | 0.56 | M250_0.6 | GGGCGGTT | 0.28 | . |
| **CTAGA**GCC | 0.33 | . | AAAAGATC | 0.56 | M148_0.6 | ATGGGCGG | 0.28 | . |
| **CTAGA**GGC | 0.34 | . | GAAAGATC | 0.6 | M250_0.6 | GGGCGGAC | 0.31 | M065_0.6 |

| Uli16 | | | Kwa14 | | | Ino17 | | |
|---|---|---|---|---|---|---|---|---|
| **8-mer** | **SVRW** | **RBP** | **8-mer** | **SVRW** | **RBP** | **8-mer** | **SVRW** | **RBP** |
| TCGAGATC | -1.63 | M348_0.6 | **ATGCC**GCC | -0.42 | . | CCCTCGAC | -0.81 | . |
| GCGAGATC | -1.55 | . | **ATGCC**GTC | -0.36 | . | TCCCTCGA | -0.77 | . |
| ACGAGATC | -1.49 | M348_0.6 | **ATGCC**GCA | -0.35 | . | CCTCGACG | -0.77 | . |
| CCGAGATC | -1.3 | . | **ATGCC**GAG | -0.35 | . | TTCCCTCG | -0.74 | . |
| TAGAGATC | -1.05 | . | **ATGCC**GCT | -0.35 | . | GTCCCTCG | -0.65 | . |
| GAGAGATC | -1.02 | M147_0.6 | **ATGCC**GGA | -0.34 | . | TAAGGCAT | -0.6 | . |
| AAGAGATC | -1.01 | . | **ATGCC**GAT | -0.34 | . | GCAGCAGC | -0.6 | . |
| ATCGAGAT | -0.94 | M348_0.6 | **ATGCC**GGG | -0.32 | . | CCCTCGGC | -0.59 | . |
| ACCGAGAT | -0.93 | . | TGCCGCCA | -0.32 | . | AGGCATTA | -0.57 | . |
| GGCGAGAT | -0.9 | . | **ATGCC**GGC | -0.31 | . | TCGAGGCA | -0.57 | . |
| CGTGATCG | 0.86 | . | GCCATGTG | 0.22 | . | CGGAACCT | 0.66 | . |
| CGAGATGG | 0.86 | . | CC**ATGCC**G | 0.22 | . | ACCGGAAC | 0.66 | M291_0.6 |
| CGTGATCA | 0.89 | . | **ATGCC**ATA | 0.22 | . | GAACCGGA | 0.69 | . |
| CGTCAGAT | 0.93 | . | **ATGCC**AAG | 0.23 | . | CCGGACCC | 0.72 | . |
| CGCGATCA | 0.94 | . | TGCCATGC | 0.23 | . | GGAACCCT | 0.72 | . |
| CGAGATCT | 0.94 | . | **ATGCC**ACG | 0.23 | . | TCGGAACC | 0.76 | . |
| AGAGATCG | 1 | . | GCCACGTG | 0.24 | . | CCGGAGCC | 0.79 | . |
| CGAGATCC | 1.01 | . | **ATGCC**AGT | 0.25 | . | GGAACCCA | 0.81 | . |
| CGAGATCA | 1.08 | . | **ATGCC**ATT | 0.25 | . | CCGGAACC | 0.92 | M291_0.6 |
| CGAGATCG | 1.11 | . | **ATGCC**ATG | 0.32 | . | CGGAACCC | 0.98 | . |

(Continue to the next page)

| HSS | | | ORI | | |
|---|---|---|---|---|---|
| 8-mer | SVRW | RBP | 8-mer | SVRW | RBP |
| GGCCGGCC | -1.68 | . | GGCCGGCC | -1.54 | . |
| CGGCCGGC | -1.39 | . | GCCGGCCG | -1.38 | . |
| GCCGGCCG | -1.37 | . | CGGCCGGC | -1.23 | . |
| GCCGGCCC | -1.24 | . | AGCCGGCC | -1.19 | . |
| GCCGGCCA | -1.2 | . | GCCGGCCC | -1.14 | . |
| AGCCGGCC | -1.2 | . | GCCGGCCA | -1.09 | . |
| GGCCGACC | -1.15 | . | GCGGCCGG | -1.03 | . |
| CGGCCGAC | -1.01 | . | CCGGCCGG | -0.98 | . |
| GCGGCCGG | -1.01 | . | CGCCGGCC | -0.97 | . |
| CCGGCCGG | -1 | . | GGCCGACC | -0.95 | . |
| CGGGTCAC | 0.56 | . | CGCAATTG | 0.45 | . |
| GGTCACGC | 0.56 | . | CCTCCATT | 0.46 | . |
| CCTCCGGG | 0.56 | . | GTCGCCAT | 0.46 | . |
| TTGTTCAC | 0.56 | . | GCCGCCAT | 0.48 | . |
| GTTCCTCC | 0.57 | . | ACGCCATT | 0.49 | . |
| CGGGACAC | 0.57 | . | CGCCATTA | 0.51 | . |
| GGGGTCAC | 0.57 | . | GACGCCAT | 0.52 | . |
| GGGTCACA | 0.59 | . | TCCGCCAT | 0.58 | . |
| TGGGTCAC | 0.6 | . | CCGCCATT | 0.68 | . |
| GGGTCACC | 0.6 | . | CGCCATTG | 0.7 | . |

**Supplemental Table S5: The ten most positive and negative 8-mers from the 11 data sets.** For each of the trained MTSA models, we identified the ten most positive and negative 8-mers. Matched RBP motif IDs from the CISBP-RNA database (FIMO with p-value $<10^{-3}$) are shown in the RBP column. Sequences matched to left and right flanking sequences are bolded and underlined, respectively.

| Data set ID | Model without flanking sequences | Model with flanking sequences | Difference in Correlation |
|---|---|---|---|
| **Mel12** | 0.46 | 0.54 | +0.08 |
| **Khe13** | 0.41 | 0.44 | +0.03 |
| **Khe13K** | 0.51 | 0.66 | +0.15 |
| **Kwa14** | 0.46 | 0.56 | +0.10 |
| **Ern16** | 0.44 | 0.49 | +0.05 |
| **Ern16K** | 0.37 | 0.56 | +0.19 |
| **Tew16** | 0.53 | 0.56 | +0.03 |
| **Uli16** | 0.63 | 0.71 | +0.08 |
| **Ino17** | 0.63 | 0.68 | +0.05 |

**Supplemental Table S6: Adding flanking sequences to tags significantly improves model performance.** MTSA models were retrained by removing flanking sequences. The correlation between the observed relative expression and the predicted expression by these models are shown in the 2$^{nd}$ column. The correlation values in the 3$^{rd}$ column are the same as the *r* values shown in **Fig. 2A** and **Supplemental Fig. S2**. Differences in the correlations are shown in the 4$^{th}$ column.

| Data set ID | Minimum DNA count per tag (-m) | Minimum number of tags per CRE (-t) | Left flanking sequence (-l) | Right flanking sequence(-r) | Number of tags in training |
|---|---|---|---|---|---|
| **Mel12** | 1,000 | 5 | CTAGA | AGATC | 12,689 |
| **Khe13/Khe13K** | 500 | 5 | CTAGA | AGATC | 25,344 |
| **Ern16/Ern16K** | 2,000 | 5 | CTAGA | AGATC | 10,127 |
| **Uli16** | 100 | 5 | CTAGA | AGATC | 63,977 |
| **Tew16** | 60 | 10 | CTAGA | AGATC | 72,435 |
| **Tew16N** | 60 | 10 | CTAGA | AGATC | 72,402 |
| **Kwa14** | N/A | 4 | ATGCC | TGAGC | 12,776 |
| **Mog13** | N/A | 4 | . | . | 2,436 |
| **Ino17** | 50 | 10 | AATTC | CATTG | 65,360 |
| **Ino17W** | 50 | 10 | AATTC | CATTG | 64,189 |
| **Kle20** | . | 10 | CTAAG | CATTG | 28,187 |

**Supplemental Table S7: List of specific parameters for building MTSA training sets.** For each of the data sets, we optimized the parameters as shown above to process and build MTSA training sets. For the Kle20 data set, we used a count per million (CPM) > 3 as a minimum DNA count.

| Data set ID | RBP | miRNA |
|---|---|---|
| **Mel12** | 47 | 0 |
| **Khe13** | 42 | 7 |
| **Khe13K** | 40 | 7 |
| **Ern16** | 44 | 5 |
| **Ern16K** | 42 | 4 |
| **Uli16** | 42 | 7 |
| **Tew16** | 81 | 20 |
| **Kwa14** | 36 | 9 |
| **Ino17** | 45 | 11 |
| **HSS/ORI** | 56 | 19 |

**Supplemental Table S8: Number of miRNA and RBP features selected for multivariate regression analyses.** For each of the data sets, we selected miRNA and RBP that are likely to affect tag expression based on their expression and match frequencies in tags.

### Supplemental Notes

While the original study (Ulirsch et al. 2016) reported 32 functional variants, we found 29 additional variants using the same data set and pipeline scripts. After consulting with the authors of the study, we discovered that the Quantile-Normalization step was not applied to the control data set in the published work. We therefore used the updated quantile normalized data. The original authors verified this issue and agreed with the usage of the updated results.
